# Transposable element small and long RNAs in aging brains and implications in Huntington’s and Parkinson’s disease

**DOI:** 10.1101/2024.10.22.619758

**Authors:** Gargi Dayama, Shruti Gupta, Brianne K. Connizzo, Adam T. Labadorf, Richard H. Myers, Nelson C. Lau

**Affiliations:** Department of Biochemistry and Cell Biology, Boston University Chobanian and Avedisian School of Medicine, Boston MA 02118; Department of Biomedical Engineering, Boston University, Boston MA 02215; Department of Neurology, Boston University Chobanian and Avedisian School of Medicine, Boston MA 02118; Professor Emeritus, Boston University Chobanian and Avedisian School of Medicine, Boston MA 02118; Genome Science Institute, Boston University Chobanian and Avedisian School of Medicine, Boston MA 02118

**Keywords:** Transposable elements, human aging, brain small RNAs, brain long RNAs, neurodegenerative diseases, biomarkers, Huntington’s disease and Parkinson’s disease

## Abstract

Transposable Elements (TEs) are implicated in aging and neurodegenerative disorders, but the impact of brain TE RNA dynamics on these phenomena is not fully understood. Therefore, we quantified TE RNA changes in aging post-mortem human and mouse brains and in the neurodegenerative disorders Huntington’s Disease (HD) and Parkinson’s Disease (PD). We tracked TE small RNAs (smRNAs) expression landscape to assess the relationship to the active processing from TE long RNAs (lnRNAs). Human brain transcriptomes from the BrainSpan Atlas displayed a significant shift of TE smRNA patterns at age 20 years, whereas aging mouse brains lacked any such marked change, despite clear shift in aging-associated mRNA levels. Human frontal cortex displayed pronounced sense TE smRNAs during aging with a negative relationship between the TE smRNAs and lnRNAs indicative of age associated regulatory effects. Our analysis revealed TE smRNAs dysregulation in HD, while PD showed a stronger impact on TE lnRNAs, potentially correlating with the early average age of death for HD relative to PD. Furthermore, TE-silencing factor TRIM28 was down-regulated only in aging human brains, possibly explaining the lack of substantial TE RNA changes in aging mouse brains. Our study suggests brain TE RNAs may serve as novel biomarkers of human brain aging and neurodegenerative disorders.

## INTRODUCTION

Transposable elements (TEs) are the most significant fraction of the human genome (∼40-50%), yet only recently have we begun to appreciate TEs’ emerging impact on human diseases from cancer to neurodegenerative disorders^1–4^. Mammalian TEs are primarily retro-elements representing the main families of LINEs, SINEs (Alu’s and SVAs), and Human Endogenous Retroviruses (HERVs)^5,6^. TE activation is readily detected in human and mouse disease models, but the significance of TEs in disease etiology remains unclear^7–10^. For example, it remains unclear whether human disease may be a consequence of TE driven genome destabilization^11–13^, or an age associated modification of TE transcription which perturbs the normal transcriptome of healthy cells^2,14,15^.

To maintain stability in complex genomes despite the massive composition of TEs^5,16^, animals evolved multiple pathways to silence TE expression, ranging from RNA interference (RNAi) with silencing small RNAs (smRNAs) to histone modification regulators like Krüppel-associated box (KRAB) zinc finger proteins (KZFPs)^17–19^ and the HUSH Complex that silences TEs and KZFPs^20,21^. Germ cells and embryonic cells are the evolutionary battlegrounds between the host animal genome and selfish TEs. However, in germ cells, highly expressed TE transcription can be offset by TE-silencing Piwi-interacting RNAs (piRNAs) and small interfering RNAs (siRNAs)^22^. In contrast, human embryos and embryonic stem cells repress TEs like HERV-K and SVAs by KZFPs such as ZNF417 and ZNF587^18,19^, while somatic cells rely additionally on the HUSH complex to maintain this silencing throughout development^23,24^.

From mammals to model organisms like nematodes and *Drosophila,* young healthy animals start out with effective TE silencing mechanisms. Yet with physiological decline during aging, multiple studies corroborate the activation of TE expression^3,25–31^. Cellular senescence is one biological phenomenon triggering TE activation and contributing to age-related inflammation and disease^3,32–34^. Perhaps the breakdown in silenced chromatin architecture during aging-associated senescence exposes TEs to promiscuous transcription or leaching of cytoplasmic DNA copies that trigger the same innate immune response that protects against virus replication^35–37^.

Therefore, TE dysregulation has emerged as a major hypothesis to explain aging-associated diseases and neurodegenerative disorders^3,9,38^. Neuronal damage from inflammation has been connected to elevated HERV-K TE expression in Amyotrophic Lateral Sclerosis (ALS)^39^, while Parkinson’s Disease (PD) progression has been associated with LINE-L1 and SVA-76 TE expression and polymorphisms^8,40^. The brain RNA-binding protein TDP-43 is implicated in various age-related diseases including Alzheimer’s Disease (AD), frontotemporal dementia (FTD), and PD^41,42^. *Drosophila* and cellular models of TDP-43 loss of function also exhibit clear TE activation^26,43–46^. TE activation in brains and neurons have also been linked to polymorphisms and pathogenic expression of the MAPT gene that generates the tau protein, a factor strongly associated with AD^10,47–50^. The neurodegenerative disorder of Huntington’s Disease (HD) has only a modest literature investigating the role of TEs in this disease, with just a handful of studies^51–54^.

Despite this progress, there is still an incomplete understanding of the biological implications of activated TE RNA expression during mammalian brain aging and in neurodegenerative disorders. To further our understanding of their role, this study performs an in-depth analysis of the relationships between TE long RNAs (lnRNAs) and TE small RNA (smRNAs) expression patterns particularly in the human and mouse brain. In this novel study, we analyze matched high-throughput sequencing profiles of both TE smRNAs and lnRNAs from comprehensive datasets associated with aging, PD and HD (**Figure 1**).

**Figure 1.**
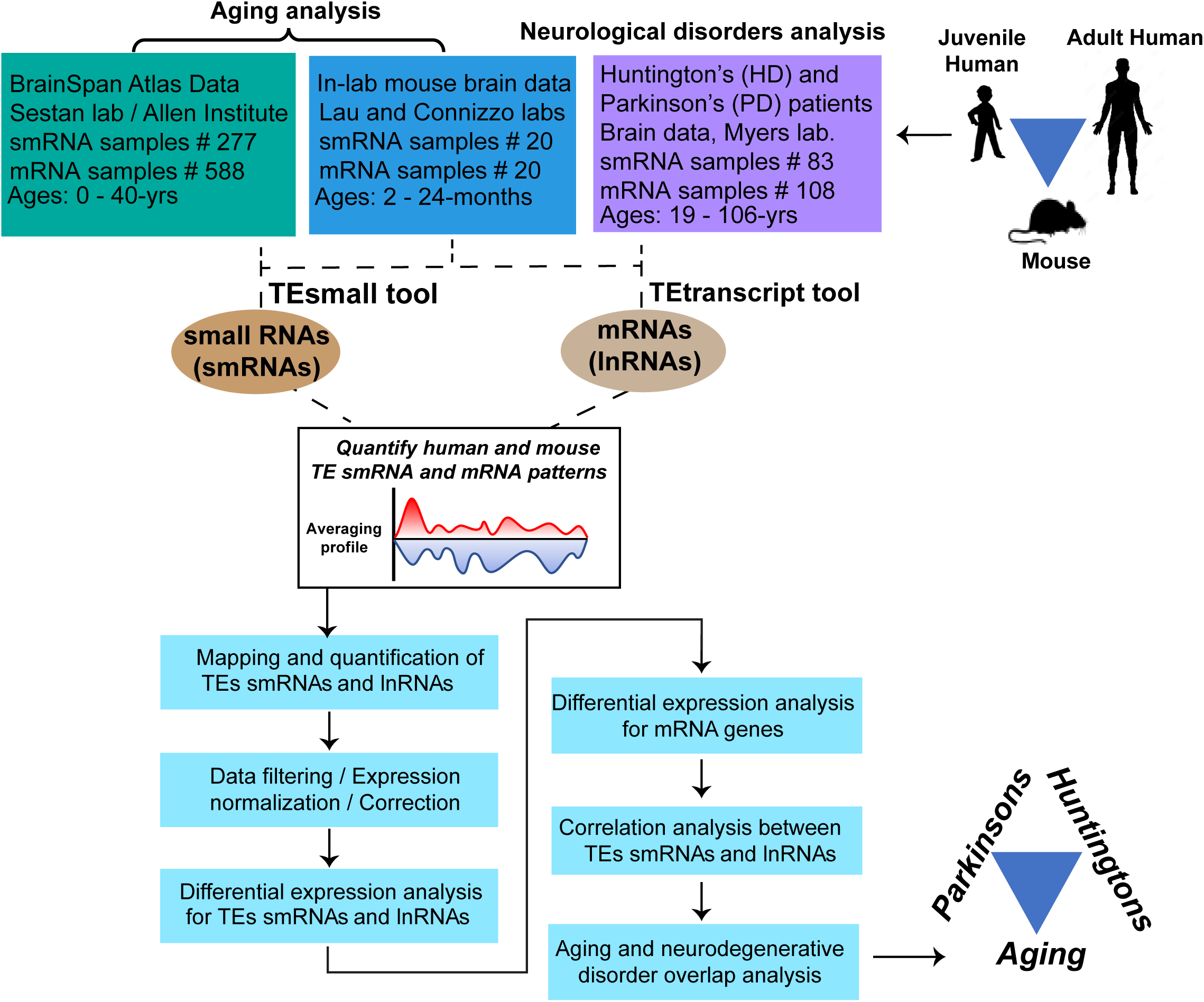
Genomics study overview. Diagram of our genomics process to investigate the connection between small RNAs (smRNAs) and mRNAs (lnRNAs) arising from Transposable Element (TE) transcripts in the human and mouse brain. Our study focuses on normal aging and in two human neurolog ical disorders, Huntington’s disease (HD) and Parkinson’s disease (PD).

Our study’s goals are to (1) determine which TEs are most frequently activated during aging, (2) show that TE lnRNA activation goes beyond the passive expression pattern of TE smRNAs, and (3) illuminate how human and mouse brains process TE lnRNAs selectively into potentially regulatory patterns of TE smRNAs. To achieve these goals, we investigated the spatiotemporal expression patterns of human TE RNAs across different brain regions and ages from childhood to adulthood (0-40 years)^55,56^ using location-specific method for quantification. Previous studies have also measured TE expression in human transcriptomes utilizing TE consensus entries to represent the numerous repeated TE copies scattered throughout the genome^10,18,19,47,50,57–61^, hence we further bolstered our results with a consensus-based quantification as well.

Our comparison of aging human and mouse brains, reveals that human brain TE RNA structure not only show a major age-related shift around the age of 20years, but also differ from mouse brains which lack TE RNA changes during aging. Finally, we performed datamining from matched smRNA and lnRNA datasets for TE RNA signatures specific to HD and PD^62–65^, and identified TE profiles as potential novel biomarkers for PD and HD.

## RESULTS

### Impact of aging on human and mouse brain TE smRNA expression patterns

To characterize TE smRNA expression in the aging brain, we analyzed 277 smRNA samples across 16 brain regions in 22 individuals from 0-yrs to 40-years of age from the BrainSpan Atlas dataset^55,56^ and also generated our own transcriptome datasets from ten 4-month old and ten 24-month old mouse brains (**Extended Data Table 1**). We first used the TEsmall pipeline^66^ on this human brain dataset. As expected, microRNAs (miRNAs) are the majority of the smRNA reads (∼30-90%), followed by structural RNAs at ∼10-40% (**Extended Data Figure 1a**). Inspection of the minor smRNA categories showed that anti-sense TE smRNAs are the next majority (1-4%), followed by sense TE smRNAs (∼0.1-0.5%) and reads annotated to piRNA cluster loci (0.001-0.006%) (Extended Data Fig. 1b).

We examined the effect of age on TE smRNA expression and found structural smRNAs, piRNAs and sense TE sRNAs were significantly increase in older brains (21-40-yrs), with overall miRNAs being significantly decreased. No change was found for anti-sense TEs between younger (0-20 yrs) and older (21-40 yrs) age groups (Extended Data Fig. 1c). Similarly in the mouse brain, the majority of the smRNAs were miRNAs, followed by structural RNAs. In contrast to humans, mouse brain sense TE smRNAs were more abundant compared to anti-sense TE and piRNA cluster smRNAs (**Extended Data Figure 2a, b**). Furthermore, none of the smRNA categories showed any significant change in abundance based on age (Extended Data Fig. 2c).

To gain insight into the overall brain TE smRNA data structure, we performed principal component analysis (PCA). Human brain TE smRNA profiles showed a clear clustering based on age, separating pre-adults (0-20 yrs) from adults (21-40 yrs), whereas mouse brain TE smRNAs did not have age-related clustering (**Figure 2a**). The PCA prompted us to define two human age groups: 0-20yrs and 21-40yrs for subsequent differential expression analysis (DEA). The same age-based clustering was evident through PCA (Extended Data Fig. 3d) when using average counts across brain regions for each of the 22 individuals, ruling out the possibility that this clustering was driven by brain region artifact. After correcting for RNA Integrity Numbers (RIN, **Extended Data Figure 3a**) and brain region biases in human and just RIN in mice, DEA was performed using the DESeq2 package^67^.

**Figure 2.**
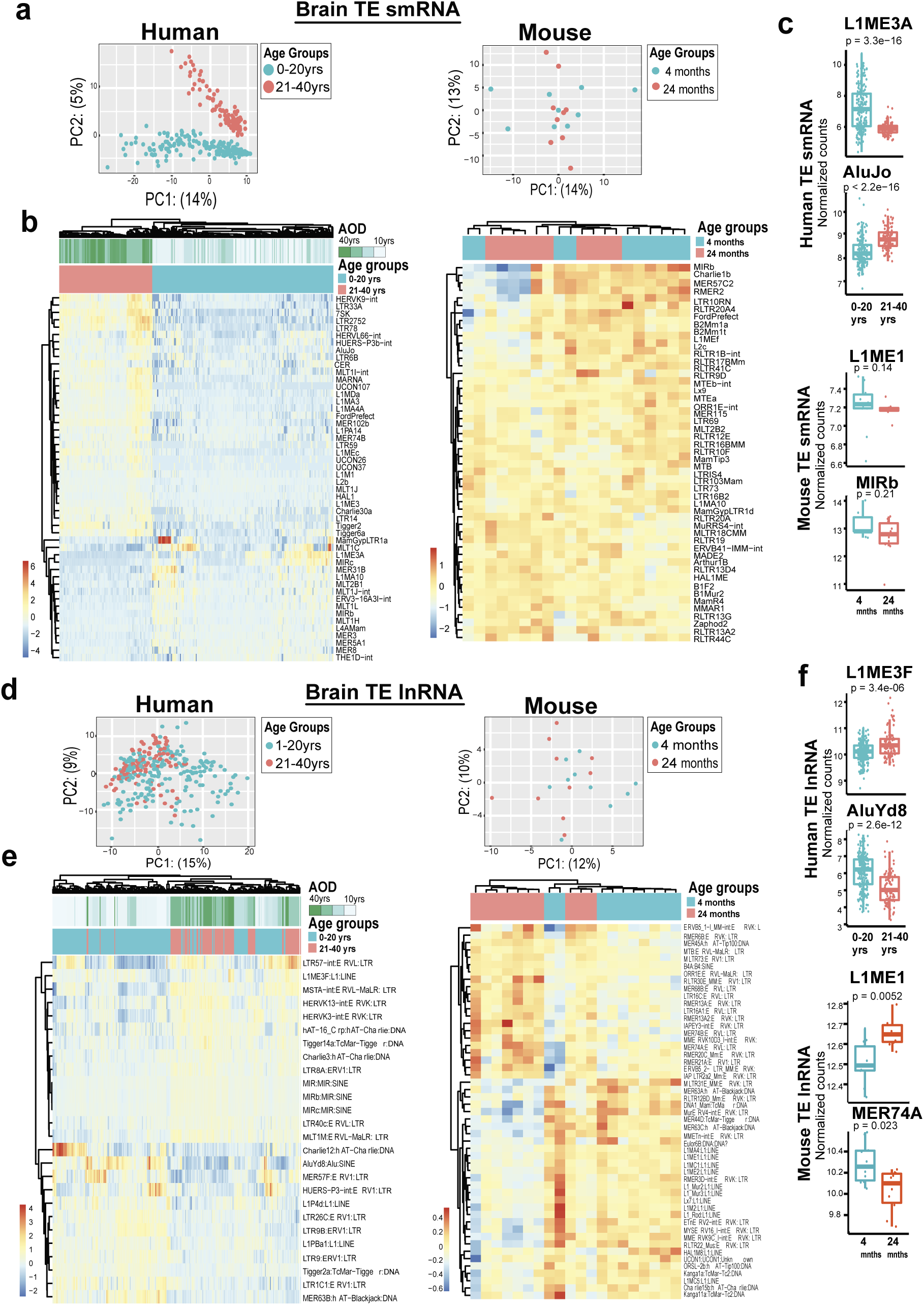
Differential expression analysis of TE smRNAs and lnRNAs in aging humans and mice brains. (**a**) Principal Component Analysis (PCA) of TE smRNA in aging human BrainSpan data showing a clear clustering between pre-adults and adults (Left), versus mouse data generated for the current study with no obvious separation based on age groups (Right). (**b**) Heatmap of hierarchical clustering of TE smRNAs per each TE family with clear clustering observed for human data but not for mouse data. (**c**) Select examples from the most significant DE smRNA TEs showing the change between the two age groups for human and mouse. (**d**) PCA of TE lnRNAs in aging human Brain-Span data and mouse brain data based on the two age groups. (**e**) Heatmap of hierarchical clustering of TE lnRNAs per each TE family for human and mouse data. (**f**) Select examples from the most significant TE lnRNAs’ DE showing the change between the two age groups for humans and mouse.

This analysis identified 259 TEs out of 1047 total repeats were significantly differentially expressed (DE) at FDR <= 0.01, in human brain TE smRNAs, of which only 35% (90 TEs) were up-regulated in older age group while 65% (169 TEs) were down-regulated (Extended Data Fig. 3e and **Extended Data Table 2**). These results were further validated by a nested analysis of variance (AOV), which controlled for potential individual variability (Extended Data Fig. 3g), bolstering our confidence in the impact of age on TE expression. In comparison, mouse brains lacked statistically significant DE in any TE smRNAs between young (4-months) and aged (24-months) samples (Extended Data Fig. 3g) based on FDR cutoff of <= 0.05. Therefore, a P-value cutoff to <= 0.05 cutoff was selected instead, where 34 (59%) TEs in the mouse brain show up-regulation of smRNAs and 24 (41%) TEs were down-regulated (Extended Data Fig. 3h and **Extended Data Table 3**).

This effect of aging in human brain TE smRNA expression is evident through hierarchal clustering heatmap of the top 50 DE TEs (Fig. 2b), with clear clustering between the two age groups (0-20yrs versus 21-40yrs). Indeed, DE of TE smRNAs are from all the categories of LINE, SINE, LTR and DNA; showing a strong effect of age across TE smRNAs irrespective of the order or class. However, the limited effect of age on TE smRNA in mouse is reflected by the absence of clustering in the mouse heatmap. We highlight two human TEs with significant TE smRNA DE: the LINE-L1ME3A (down-regulated, Wilcoxon test P-value=3.3e-16) and SINE-AluJo (up-regulated, Wilcoxon test P-value=2.2e-16, Fig. 2c). LINE-L1ME1 and SINE-MIRb are two mouse TEs with modest, insignificant (Wilcoxon test P-value > 0.1, Fig. 2c) TE smRNA changes.

### TE lnRNA expression landscape in aging brains

We next asked whether TE smRNAs share the same expression landscapes as TE lnRNAs and likely reflect degraded byproducts, or are the expression landscapes distinct and likely reflect a possible regulatory mechanism such as RNAi activity? To answer this question, we analyzed matched lnRNA datasets of 588 samples from BrainSpan Atlas with the TEtranscript pipeline^68^. We applied the same PCA and DEA described above to compare the TE lnRNAs for the 0-20 yrs and 21-40 yrs age groups. The cluster separation of TE lnRNA expression in PCA was much less distinct compared to the TE smRNA PCA, while the 0-20 yrs samples were also more dispersed, suggesting larger variability in the TE lnRNA expression landscape (Fig. 2d).

Nevertheless, of the 524 human TE lnRNAs differentially expressed at FDR <= 0.01 across BrainSpan Atlas samples, 308 (59%) were significantly down-regulated in the older 21-40 yrs age group and 216 (41%) were up-regulated (Fig. 2e, Extended Data Table 2, and Extended Data Fig. 3i). The hierarchical clustering heatmap reinforced the aging effect in distinguishing the top DE in human brain TE lnRNAS (Fig. 2e). Mouse TE lnRNAs analyzed for overall data structure using PCA also did not show distinct clustering between young 4-month and aged 24-month brains (Fig. 2d). However, DEA (FDR <= 0.01) represented through hierarchical clustering heatmap identified 15 down-regulated and 10 up-regulated TE lnRNAs in aging mouse brains (Fig. 2e, Extended Data Table 3 and Extended Data Fig. 3j). These results resonates with the biological differences separating humans from mice^69^ and the much shorter life-span of mice.

Notable examples of the DE of human brain TE lnRNAs during aging are represented by LINE-L1ME3F (up-regulated, Wilcoxon test P-value=3.4e-06) and SINE-AluYd8 (down-regulated, Wilcoxon test P-value=2.6e-12, Fig. 2f). Notable DE in mouse TE lnRNAs is exemplified by LINE-L1ME1 (Wilcoxon test P-value=0.0052) and LTR-MER74A (Wilcoxon test P-value > 0.1) which both did not show significant change in smRNAs, but its TE lnRNAs were up-regulated and down-regulated, respectively, in 24-month aged mice (Fig.2f). These data indicate variable expression of TE lnRNAs and TE smRNAs in the normal human brain which exhibits a shift in the TE structure during aging.

### Human brain regions reflect regulatory mechanisms directing distinct TE smRNA accumulation patterns

The BrainSpan Atlas provides a unique opportunity to examine region-specific variation of TE smRNAs and TE lnRNAs during development and aging in the human brain. When considering all TE smRNAs quantified by the TEsmall pipeline^66^, there is a enriched expression of sense-strand-mapping TE smRNAs in human brain cortex regions comprising the neocortex (**Figure 3a, 3b**). This systematic up-regulation of TE smRNAs in the neocortex is enhanced in aging human brains in the 21-40 yrs group.

**Figure 3.**
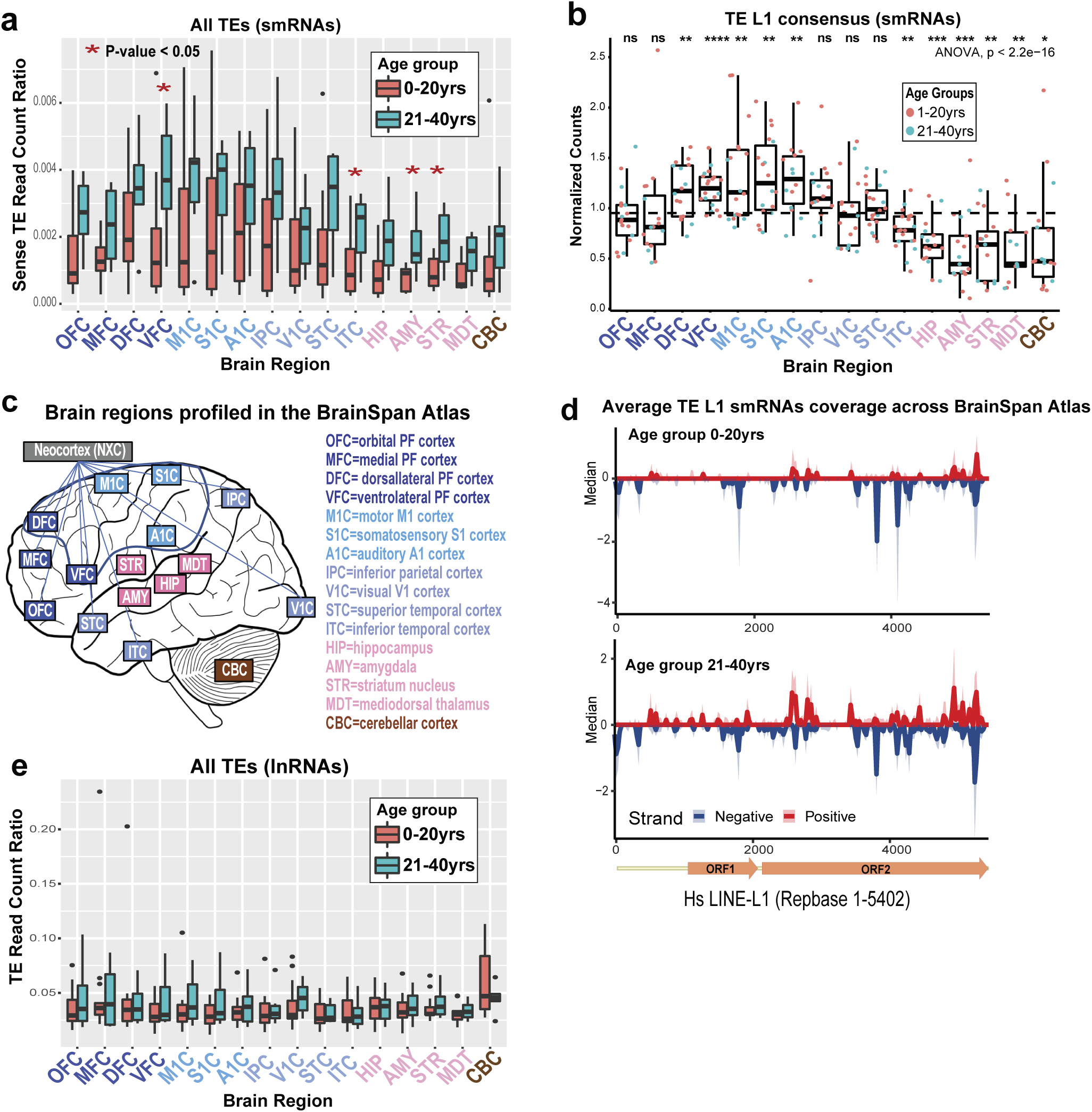
TE RNAs expression assessed across specific human brain regions. (**a**) Combined box plots quantitating all TEs expressing small RNAs across the BrainSpan regions as measured by the TEsmall pipeline. A relatively higher overall level of TE small RNA expression is observed in neocortex regions relative to other brain regions, while red asterix mark significant differences in TE small RNA expression in particular regions in aged adults versus younger humans. (**b**) The neocortex enhanced L1 consensus small RNA patterns are also observed in a distinct in-lab small RNA genomics pipeline with jitters color coded by young versus aged adult brain samples. This distinct small RNA genomics pipeline results corroborates the TEsmall pipeline result in (**a**). (**c**) Diagram of the human brain regions represented in this BrainSpan Atlas dataset. (**d**) Coverage plots showing the mean coverage (thick red and blue lines for Plus and Minus genomic strand matches) and 90% confidence interval in shaded colors of the small RNAs mapping to the LINE-L1 consensus element. TE small RNA coverage clearly increases in aged versus younger human brains. (**e**) Analagous box plot as in (A) but for the TE long RNAs mapped to the TEs witht he TEtranscript pipeline. The TE long RNA patterns contrast against the small RNA patterns suggesting TE small RNA levels are moduleated distinctly from TE long RNAs.

However, TE lnRNAs as a whole dataset did not show the same neocortex-elevated expression patterns as the TE smRNAs, regardless of the young or aged groups of BrainSpan Atlas samples (Fig. 3c). Amongst the top 50 most highly expressed TE lnRNAs plotted in a heatmap, only the cerebellum exhibited a peculiar, elevated expression of TE lnRNAs particularly in the younger 0-20yr cohort (Fig. 3c and Extended Data Fig. 4c).

To confirm the striking finding of elevated TE smRNAs in the neocortex and bolster the TEsmall pipeline results, we also applied our TE-consensus-sequence pipeline for brain TE smRNA quantification^70^. The LINE-L1 consensus TE sequence mapped smRNAs that were significantly enriched (ANOVA, P-value < 2.2e-16) in the neocortex region and down-regulated in the other brain regions (Fig. 3d), consistent with the location-specific quantification of sense TE smRNAs (Fig. 3a).

We also applied the TE consensus-sequence-based quantification method that can account for sample variability in TE smRNA expression levels to generate summary coverage plots that display the mean coverage across the two age groups from human aging data. Interestingly, an increased mean coverage was observed for TEs such as LINE-L1 (Fig. 3d) and SINE-Alu in aged adults (21-40 yrs) compared to the younger 0-20 yrs cohort (Extended Data Fig. 4a). Consistent with the observed shift in TE smRNA expression, we noted a significant difference in the coverage patterns between the negative and positive strands for the younger 0-20yr cohort in LINE-L1 smRNAs that was evident in aging adult (21-40 yrs) LINE-L1 smRNAs. The TE smRNA peaks being biased for the 3’ end of the LINE-L1 consensus element is consistent with genome bias for more truncated copies of the LINE-L1 repeats in the human genome favoring more the 3’ end of LINE-L1 sequences^61,71,72^.

### The relationship between human brain TE smRNAs and TE lnRNAs during aging

Having matched smRNA and mRNA sequencing dataset enabled us to investigate the relationship between TE smRNA and lnRNA to identify the presence of a possible regulatory mechanism. We first performed a paired correlation analysis for all annotated TE in both human and mice aging datasets. While no correlation bias was observed between smRNA-lnRNA within the mouse dataset, the human data showed that most of the SINE TEs have negative smRNA-lnRNA correlation (**Figure 4a**), suggesting RNAi activity. Furthermore, if we limit our analysis to only differentially expressed TEs during aging, the negative correlation bias becomes even more pronounced for SINE as well as LINE TEs, while LTR exhibit a bias towards positive correlation (Fig. 4a, 4b).

**Figure 4.**
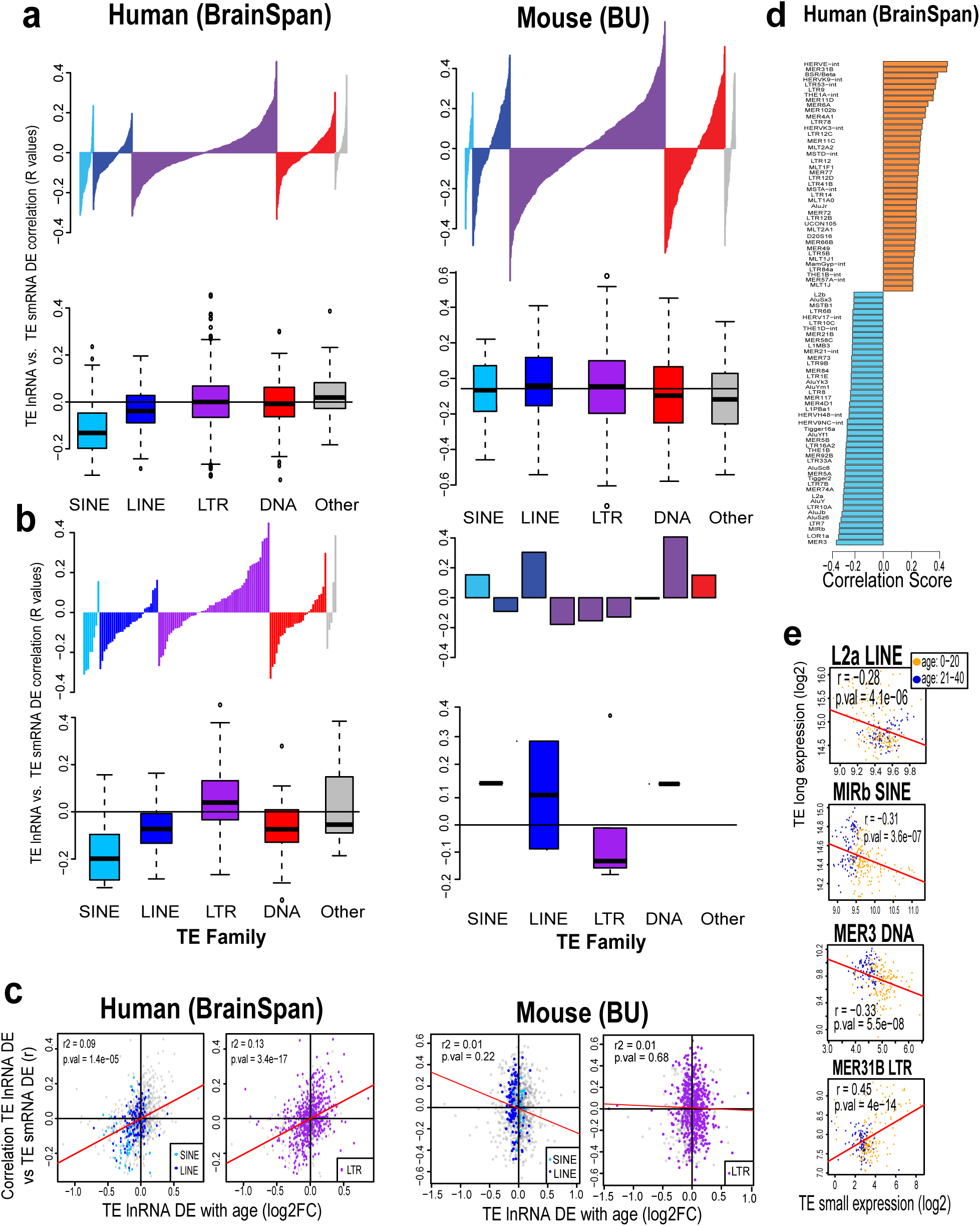
Correlation analysis of the TE long and small RNA expression during human and mouse aging brains. (**a, b**) Distribution of Pearson’s correlation score between long RNA and small RNA (**a**) for all TEs and (**b**) for shared significant DE TEs in human pre-adults and adults and young and aged mouse, color coded by TE family. (**c**) Correlation (adj. r2) between log fold change value of DE long RNA TEs and Pearson’s correlation score between TE long RNA and TE small RNA in human BrainSpan and mouse brain aging data. (**d**) Barplots displaying the most significantly correlated DE lnRNA TE and smRNA TEs in human aging data. (**e**) Scatterplot of select examples of highly correlated (Pearson’s r) TE smRNA and TE lnRNA expression in human aging data.

These results suggest that the activation or repression of several TE lnRNA during aging in human are correlated to their corresponding TE smRNAs. Consequently, we compared the magnitude of lnRNA expression change (Log fold change) during aging to the smRNA-lnRNA Pearson’s correlation score (Fig. 4c). Interestingly, we observed a highly significant positive relationship between the two metrics when considering either SINE/LINE (P-value = 1.4e-05) or LTR (P-value = 3.4e-17) TEs. This indicate that TEs for which lnRNAs are activated during aging also shows accumulation of smRNA, while repressed lnRNA TEs showed mostly negative smRNA-lnRNA correlation and are potentially under RNAi-like regulation. No such trends were observed in our mouse brain dataset, which can be linked to the absence of aging associated TE smRNA.

Finally, we focused on the TEs with high smRNA-lnRNA correlation (Pearson’s *r* >= 0.2) and found prominent LINE TEs (e.g.: L2a, L2b, L1PBa1, L1MB3) and SINE TEs (e.g: AluJb, AluY, AluSz6, AluSx3) with negative correlation (Fig. 4d). In contrast, TEs with positive smRNA-lnRNA correlation were mostly from LTR subgroup of HERVs and MERs. The scatterplots of smRNA-against lnRNA-abundance allows us to fully appreciate the connection between the two forms of RNA and their behavior relative to aging. We here highlighted four examples with LINE-L2a (P-value = 4.1e-06) and LTR-MER31b (P-value = 4e-14), both down-regulated during aging, but have inverted relationships with smRNA, while DNA-MER3 (P-value = 5.5e-08) and SINE-MIRb (P-value = 3.6e-07) both shows negative lnRNA-smRNA correlation and upregulation of lnRNAs during aging (Fig.4e). Overall, this analysis suggests that the relationship between smRNA and lnRNA has a strong influence over the TE expression during aging. For instance, LINE/SINE TEs, mostly repressed during aging, are under negative regulation with their corresponding smRNAs.

### Impact of PD and HD on human brain TE smRNAs and TE lnRNAs

We also extended our analysis to the neurological disorders with published brain transcriptome datasets containing extensive matched lnRNA and smRNA profiles. These were human frontal cortex total RNAs from healthy control individuals and patients with diagnosed Parkinson’s disease (PD) and Huntington’s disease (HD) generated by the Myers lab^62–64,73^(**Extended Data Table 4**).

To determine whether PD and HD impact human brain TE smRNA and TE lnRNA levels, we extended our analysis first to 83 smRNA libraries from two cohorts (1st cohort: PD n=28, Control n=11; 2nd cohort: HD n=21, Control n=23). The overall smRNA annotation profiles in this PD-HD-Control dataset were similar to the BrainSpan Atlas datasets: miRNAs dominated followed by structural smRNAs (**Extended Data Figure 5a, b**). Anti-sense TE smRNAs (∼1-3%) were more abundant than sense TEs (∼ 0.1-0-4%), and annotated piRNA cluster loci reads were least prevelant (∼ 0.001-0.005%). When comparing the PD and HD smRNA functional annotations to healthy controls, the two disorders also displayed clear molecular differences, such as elevated miRNAs in HD and lower structural smRNAs in HD, and vice versa in PD (P-value <0.05) (**Extended Data Figure 6a, 6c**).

To support the assessment, whether human brain TE smRNAs are processed by regulated biological processes and are not just byproducts of TE lnRNA turnover, we first noted that the TE smRNAs coverage patterns are not random and come from both plus and minus strands (Fig. 3d). Furthermore, we tracked the TE smRNA length distributions and sequence logos from HD and PD datasets (Extended Data Fig. 7a, 7b), and found these brain TE smRNA lengths (peaking between 23-30nt) and sequence logos (a bias for U/T as the first 5’ nucleotide) to be similar to piRNAs that are typically most highly expressed in the testes and are generated by the piRNA pathway of enzymes to target TE-specific silencing in gonads^74,75^.

Focusing on TE smRNAs, PCA did not show separation in PD brains versus controls, whereas HD brains showed a striking dispersion of TE smRNA signatures that possibly reflect a spectrum of HD severity (**Figure 5a, 5b**) compared to controls that were clustered together. After correcting for batch effects, our DEA results of TE smRNA changes were consistent with the PCA: there were 5 times more TE smRNAs showing DE (FDR <= 0.01) in HD to control versus PD to control (FDR <= 0.05) (**Extended Data Table 5 and Extended Data Fig. 6b, 6d)**. Furthermore, hierarchical clustering analysis of significantly DE TE smRNAs was much more effective at clustering HD samples together than PD samples (Fig. 5a, 5b). Some specific TEs like LINE-L1MD1 (P-value = 0.017) and LTR-HERVL-int (P=value = 0.00019) down-regulated TE smRNAs in PD compared to controls, but overall the TE smRNA changes were more prominent for HD, as represented by TE smRNA decreases in SINE-AluJb, (P-value = 0.00031) and LTR-HERVK-int (P-value = 0.00063) (Fig. 5c).

**Figure 5.**
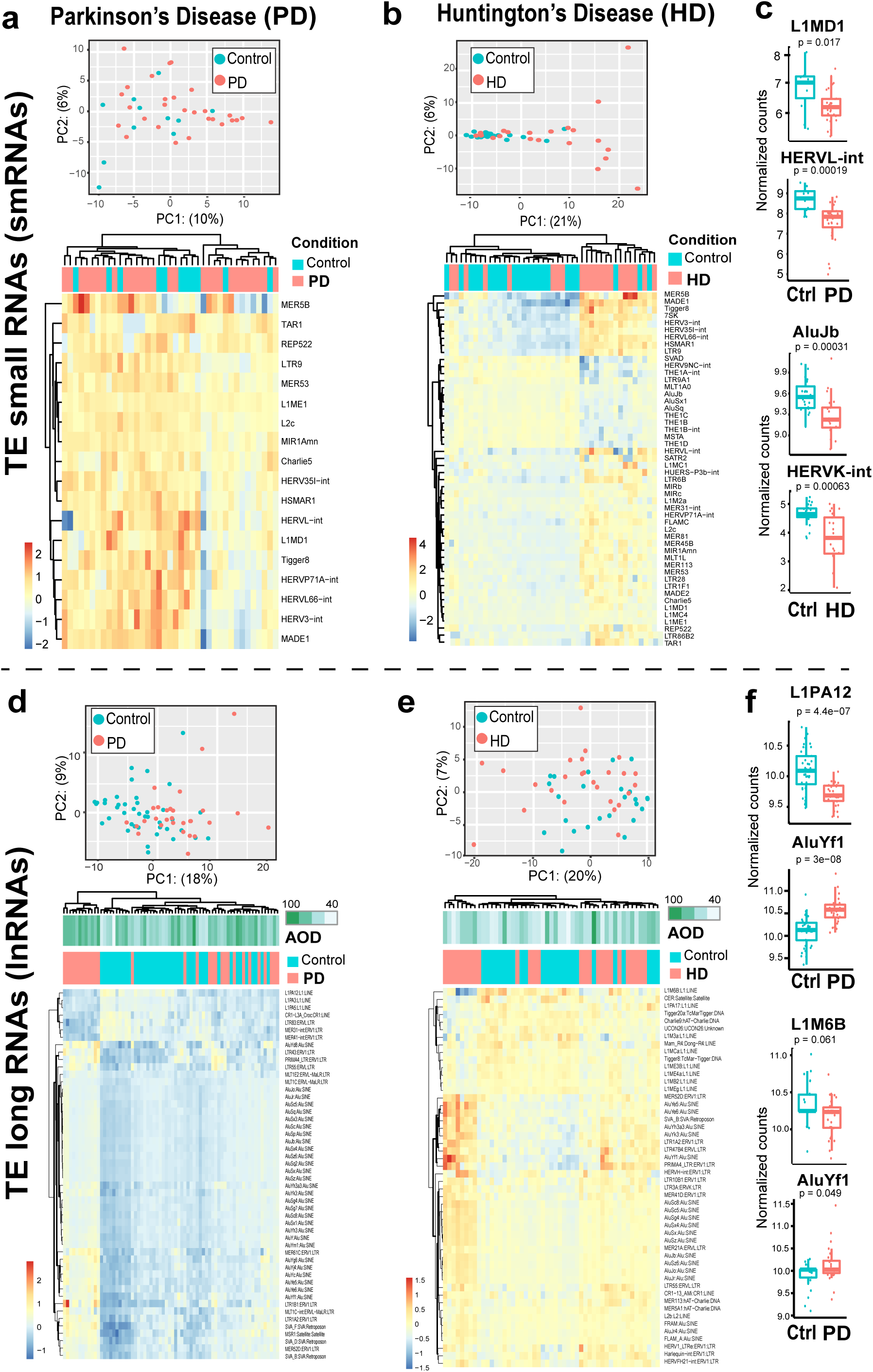
Expression patterns of TE transcripts in brains with Parkinson’s Disease (PD) and Huntington’s Disease (HD). (**a,b** and **d,e**) Top sections are Principal Component Analyses (PCAs) of TE smRNA and lnRNA expression patterns in brain samples afflicted with PD (**a,d**) and HD (**b,e**). Lower sections below PCAs are Hierarchical Clustering Heatmaps of the TE smRNA and lnRNA expression patterns. Age of death (AOD) was only correlated with the clustering patterns of the TE lnRNA expression patterns with PD as noted in (**d**, Heatmap). TE examples each in PD and HD brains where TE smRNA expression (**c**) and TE lnRNA expression (**f**) are significantly affected in the neurodegenerated state. All p-values are from Wilcoxon tests.

After applying a similar batch-correction as above (TE smRNA) to TE lnRNAs from PD and HD datasets, PCA and DEA (FDR <= 0.01) analysis showed clearer delineation of TE lnRNAs in the PD-control cohort compared to the HD-control cohort (Fig. 5d, 5e). The PD-control cohort displayed 285 TEs with DE of TE lnRNAs versus just 56 TEs with DE of TE lnRNAs in the HD-control cohort (**Extended Data Table 5** and **Extended Data Figure 9A, 9B).** Amongst the top 50 DE TE lnRNAs, a group of PD samples were distinctly clustered and consistently had higher Age of Death (AOD) potentially suggesting an increase in disease severity with age, also confirmed by increasing BRAAK score with AOD (Fig. 5d and **Extended Data Figure 8c**). Furthermore, many Alu sub-family TEs from SINEs were up-regulated in PD and showed a strong co-regulation across samples. The HD cohort’s TE lnRNA dataset were slightly more variable, and even after removal of some outliers (Extended Data Fig. 8d, 8f), this analysis still lacked a clustering group based on TE lnRNAs expression and AOD signature (Fig. 5e). Boxplots for representative TEs (Fig. 5f) reflect how the TE lnRNAs DE is much greater in the PD cohort than the HD cohort.

### PD and HD differ in the relationships between TE smRNA and TE lnRNA expression patterns

To determine how PD and HD affect the relationship between TE smRNA and TE lnRNAs, we conducted a paired correlation analysis between all annotated TE smRNAs and TE lnRNAs similar to the BrainSpan Atlas aging data. While we had observed a negative correlation for SINEs and LINEs with aging dataset, neither PD nor HD data showed any prominent correlation bias when looking at all TEs irrespective of their relationship to the disease status (**Figure 6a**). However, limiting the analysis to only DE TEs in PD, a modest positive correlation bias was observed (Fig. 6b) for SINE and LTR.

**Figure 6.**
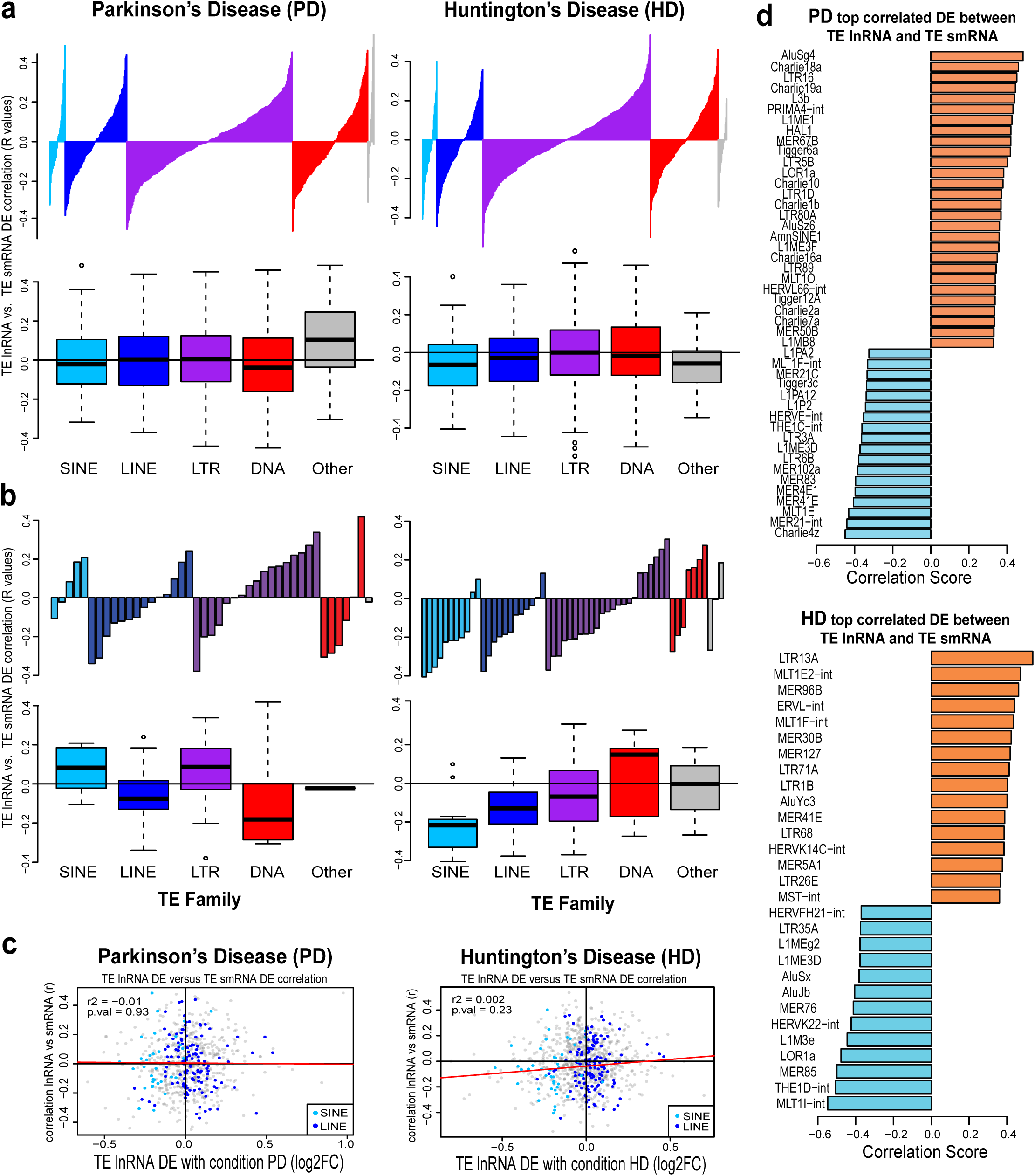
Correlation analysis of the TE long and small RNA expression in normal aging human brains and brains from PD and HD patients. (**a, b**) Distribution of Pearson’s correlation score between long RNA and small RNA (**a**) for all TEs and (**b**) for shared significant DE TEs in PD and HD, color coded by TE family. (**c**) Correlation (adj. r2) between log fold change value of DE long RNA TEs and Pearson’s correlation score between TE long RNA and TE small RNA in Parkinson’s disease (PD) and Huntington’s disease (HD). (**d**) Barplots displaying the most significantly correlated DE lnRNA TE and smRNA TEs.

In contrast, we saw a striking negative correlation bias for DE LINE and SINE TEs in HD similar to the one observed in the BrainSpan Atlas cohorts (Fig. 6b). Noting that the comparisons in HD and PD to controls is constrained by the fact that all of PD human samples are already skewed towards aged samples, the HD cohort’s TE-smRNA/TE-lnRNA relationship result may suggest HD exacerbates the influence of normal brain aging on TE RNA regulation.

In scatterplots of Pearson’s correlation score for TE-smRNA/TE-lnRNA compared to the magnitude of lnRNA expression change (Log fold change) between the disease and control samples, this analysis masks the trend of distinctly correlated regulatory changes across all the TE RNAs in both the PD and HD datasets (Fig. 6c). Perhaps the paired correlation analysis could not establish statistically significant correlations (PD: P-value = 0.93 and HD: P-value = 0.23) with the entirety of the different human TE families because the constraint of these cohorts all being skewed towards aged brains, and the diversity of TE families not all subjected equally to the impact of PD and HD etiology.

Specific TEs in PD and HD cohorts that showed either a strong positive or a negative correlation (Pearson’s *r*^2^ >= 0.3) are highlighted in the Fig. 6d bar plots. The main notable observation is that the directionality of the correlation scores for the SINE-Alu family (AluJb, AluYc3, AluSx, AluSz6) in the HD cohort is consistent the correlation score directionality in the BrainSpan Atlas aging cohorts (Fig. 4d). With the younger onset of HD compared to PD (Extended Data Fig. 8a), the possible dysregulation of TE RNAs that we observe in normal aging could be reflected in HD as well by our correlation analysis.

### Integrating the DE patterns of genes and TEs RNAs to define potential TE biomarkers for tracking human aging and neurodegenerative disorders

Although TE sequences make up the bulk of the human genome’s DNA, the functional readout of the genome has been primarily attributed to the protein coding genes. Thus, also analyyzed the interplay between TE RNAs and protein coding genes. our study’s holistic approach was more pragmatic to be able to integrate gene regulation with TE regulation across the 1056 human brain samples in this study (Fig. 1).

To integrate the analysis of gene regulation and TE changes within the BrainSpan Atlas and the PD and HD datasets, we first conducted differential expression analysis by utilizing the gene counts from TEtranscript pipeline. Unlike TEs, for both human and mouse aging lnRNA data, through DEA (FDR <= 0.01) the effect of age on the gene expression was very evident. We observed distinct clusters of top DEGs for the two age groups (0-20yrs and 21-40yrs for human and 4month and 24month for mouse) (Extended Data Fig. 3k, 4d, 4e).

For the disease and control cohorts, all the datasets from the PD, HD and Control were analyzed using PCA. Some distinct clustering could delineate the different groups (Extended Data Fig. 9c), but the isolated cohorts PCA suggest HD gene transcriptomes can be delineated better from Control compared to PD gene transcriptomes from Control (Extended Data Fig. 9e, 9f). Volcano plots of the Differentially Expressed Genes (DEG) were distinct between HD and PD with similar numbers of DEGs (4624 and 4292 genes at FDR <0.01, respectively, Extended Data Fig. 9g, 9h). These baseline analyses suggest the DEG for HD is slightly more informative than the DEG for PD based on this overall data structure.

To identify which TEs have TE lnRNA DE patterns that correlate best with gene DE patterns from the BrainSpan Atlas and HD and PD datasets, we generated a network plot showing 261 significantly correlated (Pearson’s r > 0.5) common gene-TE pairs (**Figure 7a**). Two sets of human TE families stood out with clusters of correlated DEGs: the LTR-MER31-int, and multiple SINE-Alu elements, specifically AluYF1, AluYH3A3 AluYE5, AluYC and AluSC8. As the latest Telomere-to-telomere (T2T) human genome assembly revealed, AluY’s contribute significantly to transcriptional signals; and despite being extremely compact in the TE size, SINE-Alu’s are ∼10% of the human T2T genome^61^.

**Figure 7.**
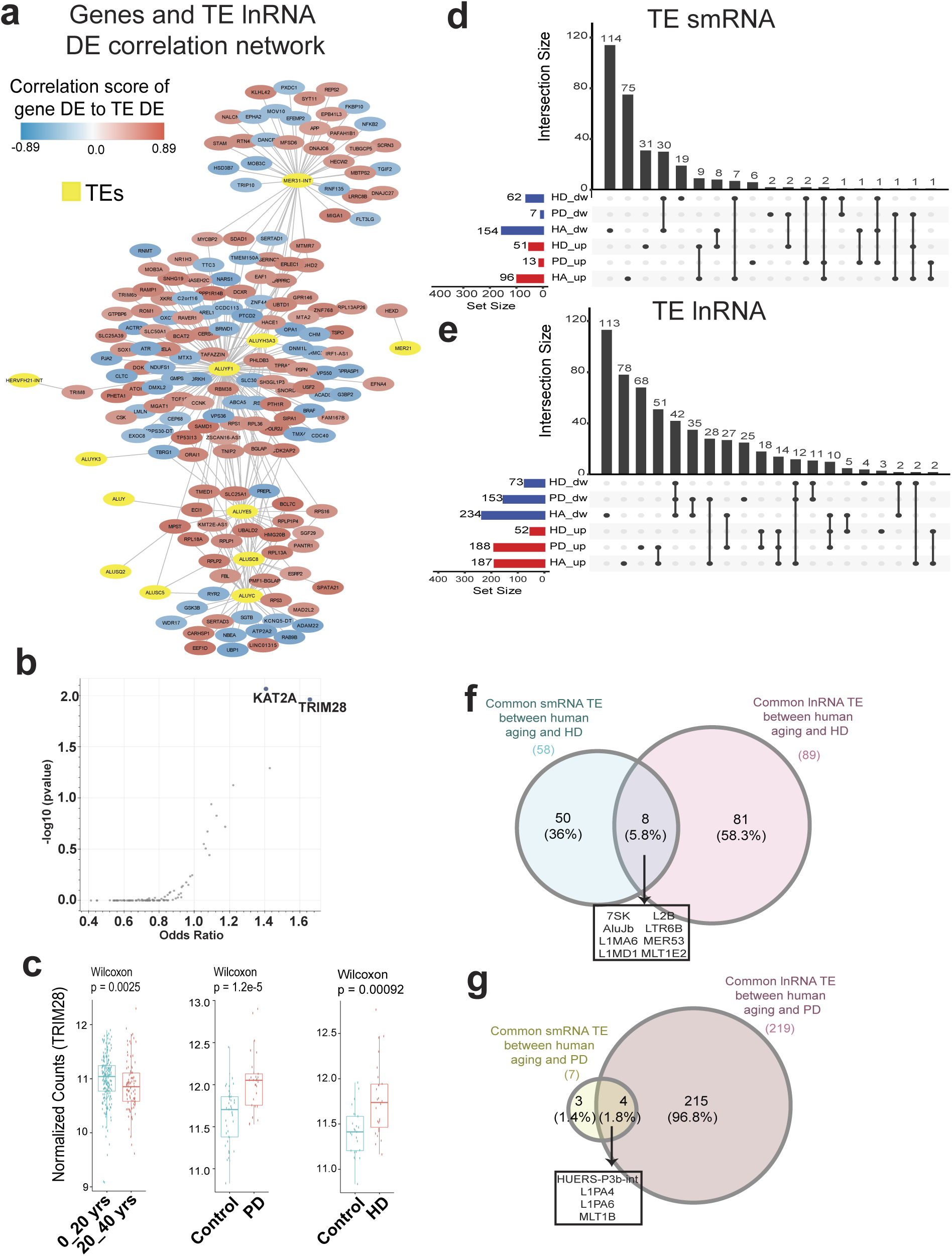
Integrative analysis of differential expression (DE) patterns in TE smRNA and lnRNA during aging and within neurogenerative disorders. (**a**) Schematic representation of correlation network for DE genes and TE long RNAs. (**b**) Enriched targets of transcription factor in significantly down-regulated genes between pre-adult (0-20yrs) and adult (21-40 yrs) with FDR cutoff of 0.05. (**c**) Boxplots with relative abundance for TRIM28 gene, between pre-adult (0-20yrs) and adult (21-40 yrs), HD and control and PD and control. (**d-e**) Upset plots linking candidate human TE transcripts as biomarkers for brain aging and neurodegeneration, based on (**d**) TE small RNAs and (**e**) TE long RNAs. (**f-g**) Venn diagrams highlighting TEs in the overlap region of shared DE patterns in TE smRNAs and lnRNAs between aging progression and Huntington’s Disease (**f**) and Parkinson’s Disease (**g**).

We mined this gene network for informative pathway terms, and both the BioCarta database^76^ and KEGG database^77^ pointed to Alzheimer’s disease (AD), ATM signaling pathways, and other cell cycle related pathways as significantly enriched terms (P-value < 0.01, **Extended Data Figure 10a, 10b**). In addition, the MER31- and AluYc-correlated gene networks connected to APP and GSK3B, which have been implicated in AD-related disorders^78–81^.

The most notable gene-to-TE-correlation result was a negative correlation of expressed TEs in aging BrainSpan Atlas adult brains and down-regulation of the TRIM28 transcription factor (TF) RNA (Fig. 7b), the only TF enriched amongst the list of annotated human TFs. Previous studies point to TRIM28 as a regulator of TEs and KZFPs during human brain development^18,19,82^. Healthy aging adult brains display lower TRIM28 levels (Wilcoxon test P-value = 0.0025, Fig. 7c), which may also implicate lower levels of protein tau^83,84^ in addition to the increased TE lnRNA levels. However, both HD and PD brains compared to controls show elevated TRIM28 (Wilcoxon test P-value = 1.2e-05 (PD) and P-value = 0.00092 (HD), Fig. 7c) that may contribute to TE and KZFP dysregulation. Lastly, we do not observe any changes to mouse TRIM28 expression between young and aged mouse brains, coinciding with the lack to TE RNA changes during mouse brain aging (Fig. 2).

To possibly define brain TE RNAs themselves as biomarkers representing normal human aging decline by connecting to dysregulation of TE levels in PD and HD, we performed intersection analyses across all three datasets, one for DE directionality of TE smRNAs and one for TE lnRNAs (DE at FDR < 0.1, Fig. 7d, 7e). HD versus control cohorts shared the most TE smRNAs with DE changes as the BrainSpan Atlas young versus aging cohorts (30 and 9 intersecting TEs, Fig. 7d). Only 9 TE smRNAs DE changes intersected between PD and HD, mostly with inverted directionality. Overall, there were only 4 significant TE smRNAs (L1MD1, L1PA4, HERVL-int, HUERS-P3b-int) with intersecting DE across three datasets irrespective of their directionality.

Many more TE lnRNAs with significant DE changes were represented in the UpSet plot of the intersecting TEs in the BrainSpan aging brains versus PD and HD (Fig. 7e). This greater number of TE lnRNAs intersections over TE smRNA intersections can be expected since sequencing coverage of TE lnRNAs is better than TE smRNAs coverage in the human brain. Nevertheless, we could further intersect these shared TE smRNAs and TE lnRNAs with DE changes between HD-Control and BrainSpan-Young-vs-Aged datasets; and PD-control and BrainSpan-Young-vs-Aged datasets in the Venn diagrams of Fig. 7f and 7g, respectively. This analysis found 8 contrasting expression patterns of TE smRNAs and TE lnRNAs suggestive of regulation during aging brains and HD-afflicted brains, including AluJb, L1MA6, L2b. The similar analysis applied to aging BrainSpan samples and PD-afflicted brains identified 4 shared TE pairs: HUERS-P3b-int, L1PA4, L1PA6, MLT1B.

From the entirety of the cross-referencing analyses in Fig. 7, we propose these candidate TE targets as potential biomarkers of human aging brains: AluYd8, AluYe6, HERVK and L1ME1. Potential TE biomarkers for studying HD could be: AluJb, L1MD1, MER53, AluY and HERVK. For PD, we propose L1PA4, L1MD1, HERVL and L1PA12 as potential TE biomarkers. Further investigations will test how well these candidate TEs can perform as potential biomarkers for assessing the prognosis of these neurodegenerative diseases.

## DISCUSSION

Our TE RNA genomics study thoroughly characterizes the dynamics of brain TE RNAs during natural human aging and in the neurodegenerative disorders of HD and PD. Importantly, our analyses rely on matched lnRNA and smRNA transcriptomes from human and mouse brains to enable correlation analyses that can provide insight into possible molecular regulatory mechanisms like RNAi. In gonads, a clear high natural expression of TE lnRNAs has been shown, that are then silenced by the Piwi proteins for which many (but not all) piRNAs are also TE smRNAs ^74^. Our analysis of the TE smRNAs from human and mouse brains suggest they share molecular characteristics of length and a 5’ U/T first nucleotide bias indicative of piRNAs (Extended Data Fig. 7) and extends previous studies also detecting TE smRNAs in mammalian brains^60,85,86^. However, since we observed human brain TE smRNA patterns change most dramatically at 20 yrs old with the onset of adulthood (Fig. 2), we wonder if the other human aging milestones of 44 yrs and 60 yrs implicated in a recent multi-omics study ^87^ may also exhibit brain TE RNA dynamics or it might just be due to a post-teen shift in growth and development in humans.

Although mouse brains express TE lnRNAs and TE smRNAs (Fig. 2), we could not detect TE RNA changes in aging mouse brains like the clear TE RNAs dynamics in aging human brains. The old-aged mouse brain transcriptomes did clearly show DE of gene’s lnRNAs (Extended Data Fig. 4e), indicating our transcriptome profiling of gene lnRNAs in mouse brains is technically sound (Extended Data Fig. 2). Two considerations we rationalize to explain the lack of TE RNA dynamics in the aging mouse brain could be: (1) our study only profiled 4 month-young and 24-month-aged male mice of the common lab inbred C57/BL6 line, so our mouse study lacks genetic diversity and may differ in aging profiles compared to the human studies; (2) the lab mice are raised in a controlled environment and exhibit biological conditions inherently different from human biology and the human condition that influences human brain development. Regarding aging profiles, one study did detect mouse TE RNA elevation in aging mouse tissues only at 36 months, but not 24 months^30^, which poses a challenge to equate a 36-month mouse equivalent to human centenarians that are not prevalent in our human transcriptome datasets. Perhaps the TE RNA dynamics differences between mouse versus human may also stem from the TRIM28 TF only decreasing in expression in human brains (Extended Data Fig. 10).

Despite both being neurodegenerative disorders, PD and HD have distinct symptoms, distinct causative gene mutations, environmental triggers, and different disease onsets: ∼30-40yrs in HD versus >60yrs in PD^88^. We are able to conclude from this data that TE lnRNAs expression changes are impacted by PD whereas TE smRNAs expression changes are a characteristic of HD in the frontal cortex of these individual brains. To help better understand what etiologies distinguish PD and HD, the measurement of TE RNAs adds a new information component towards addressing this knowledge gap. A potential interpretation is that with the older onset of PD, overall chromatin silencing architecture maintenance in those diseased brains falters more to unleash more TE lnRNAs, whereas in HD the underlying RNA processing steps to convert TE lnRNAs into TE smRNAs is being dysregulated.

Our holistic transcriptomics approach is only possible through the matched lnRNA and smRNA datasets that not only reveal the TE RNA dynamics during aging in human brains, but also the directionalities of TE RNA DE changes with correlation and network analyses. This enables us to identify specific human TE families whose RNAs may potentially serve as biomarkers for normal aging brains and neurodegenerative diseased brains. Indeed, our study indicates the TE smRNA changes that are more pronounced in HD with a strong negative correlation (SINEs and LINEs) between TE smRNA and lnRNA, aligns better with TE smRNA changes in aging brains from BrainSpan. Surprisingly, in age-related disease such as Parkinson’s, SINEs and LTRs in fact showed a positive relationship between DE smRNA and lnRNA TEs, indicating a dysregulation of TEs thus altering the relationship between the small and long RNA TEs.

However, since our input transcriptome datasets of BrainSpan Atlas^55,56^ and HD and PD cohorts^62–64,73^ were generated many years ago and the original RNAs are no longer accessible, molecular validation efforts of these candidate TE RNAs as biomarkers will require future studies with new human brain RNA samples. Nevertheless, our current study lays the foundation to next explore TE RNA regulation in other aging-related brain diseases such as AD and related dementias (ADRD) in human brains as well as mouse models of AD like the 5XFAD mouse^89^. We can then test whether the human TEs with dynamic TE RNA changes during aging and in HD and PD situations may also connect to AD and ADRD disorders. The etiology of HD and PD as motor-function-impacting disorders is clearly distinct from the memory and cognition impacts of AD and ADRD^88^, so we anticipate there to be differences in the TE RNA profiles in AD.

Indeed, some TE lnRNA expression is influenced by host gene expression when certain TE sequences reside in gene introns. Although some studies are able to attribute TE RNA impact at the locus level in cis to individual genes^90,91^. Our study identified TRIM28 transcription factor (TF), a key regulator of TEs during brain development ^82^ to be significantly down-regulated with age in human data. Interestingly, TRIM28 gene expression was up-regulated in both the neurodegenerative disorders; HD and PD compared to the control samples, which might lead to neuronal damage and inflammation ^83^. Potentially, suggesting disruption of the ‘normal’ pattern of TRIM28 as seen in healthy aging population. This finding aligns with a previous study that found the levels of pathogenic protein tau, a key player in neurodegenerative diseases, to be decreased when TRIM28 was suppressed ^83,92^.

Systems biology and genomic studies have previously overlooked TE RNAs as a contributor to aging-associated maladies because of the primacy and standardized pipelines mainly geared to protein-coding genes. Our study here helps address this limitation and enables future research efforts to unveil new potential avenues into interventions to combat the TE hypothesis of human aging and brain health decline ^3,4,14^.

## MATERIALS AND METHODS

### Human brain transcriptome datasets

The human BrainSpan Atlas dataset was generated by Allen Institute of Brain Science (www.brainspan.org, Extended Data Table 1,^55,56^) and was downloaded onto our Boston University Shared Computer Cluster onto specific securely-protected servers via the NIH dbGaP server through the approved project #18737: "Transposon expression studies in the brain." The Huntington’s disease (HD), Parkinson’s disease (PD) and neuropathologically normal control small RNA and poly-A enriched mRNA, was previously generated by the Myers lab (Extended Data Table 2,^62,64,73^). The BrainSpan Atlas data was downloaded from SRA database (BioProject ID # PRJNA242448) and the HD/PD data was also downloaded from SRA database (Bioproject ID # PRJNA271929 for RNA-Seq; Bioproject ID # PRJNA295431 miRNA-Seq). We processed ∼6M to 24M reads, with an average phred score above 30.

### Mouse brain tissue acquisition, RNA extraction, library preparation and sequencing

Mouse brain tissue of 4 months (n=10) and 24 months (n=10) old lab C57BL/6J male mice were provided by the Connizzo lab at Boston University (Extended Data Table 1). The mice had free access to food and water and were on a 12h/12h light-dark cycle. At the end of the experiment, the mice were euthanized, and their full brain dissected and flash-frozen in liquid nitrogen.

Total RNA was extracted from dissected mouse brains stored at −80⁰C. 20-50mg of frozen brain tissue was sliced off into a tube and covered with DNA/RNA Protection Reagent (NEB). Zirconium beads (3.0mm) in a bead beater were used to homogenize tissue before proceeding with NEB Monarch Total RNA Miniprep Kit (NEB #T2010), following manufacturer’s protocol. The on-column DNase I treatment was performed during RNA extraction. RNA was checked on Agilent Bioanalyzer 2100 using RNA 6000 Nano kit to get RNA Integrity Numbers (RINs) and ascertain yield.

Small RNA libraries were made using NEBNext Small RNA Library Prep Set (NEB #E7330) with up to 500ng of RNA input, following manufacturer’s protocol. Total RNA libraries were made using Zymo-Seq RiboFree Total RNA Library Kit using up to 500ng RNA input and following manufacturer’s protocol. All libraries were checked on an Agilent Bioanalyzer 2100 using either DNA 1000 or High Sensitivity DNA kit and sequenced at the BUMC Microarray and Sequencing Resource on an Illumina Next Seq 2000 using P3 flow cells for 50SE and 50PE reads for small RNA and total RNA libraries, respectively. The data can be accessed through the SRA portal BioProject ID # PRJNA1113634.

### TE smRNA and TE lnRNA data annotation, quantification, pre-processing and differential expression analysis

Small RNA data for aging (BrainSpan Atlas) and disease (HD/PD/Control) were first assessed for quality using *fastqc* then filtered for average phred score below 30 and had read length of 50bp. Adapters were trimmed using *Trimmomatic* (V - 0.39;^93^) and *TEsmall* pipeline^66^ was used for annotation and quantification of the smRNA, utilizing human reference genome build GRCh38 and mouse genome reference mm10. Relative read count breakdown of various smRNA categories were depicted using barplot in *R* (Extended Data Figs. 1a, 2a, and 5a). The TE read count were extracted using *R* script and collapsed at the sub-family level for further analyses.

Long RNA data as well was first assessed for quality using *fastqc* and trimmed for adapters using *Trimmomatic* (V - 0.39). The human and mouse datasets were annotated against human reference genome build GRCh38 and mice reference mm10, respectively, using HiSat2^94^. The SAM format file was then sorted, converted to BAM format and indexed all using *Samtools*^95^. The BAM file was fed to *TEtranscript* pipeline^68^ for quantification of TEs and gene features.

Differential expression analysis of TE smRNAs was performed using *DESeq2* package^67^ in *R*, with covariate correction informed by principal component analysis (PCA). Both the human and mouse aging datasets were corrected for RNA integrity bias (RIN, Extended Data Figs. 3a, 3f). While in humans, cause of death (COD) also showed some effect on the data structure, particularly in age group 0-20 (Extended Data Fig. 3b), it was not added as a covariate to our model due to its correlation with age which would over-correct the data (Extended Data Fig. 3c). Additionally, since the samples in BrainSpan dataset encompassed various brain regions, this variable was added as covariate as well. To further validate our findings and ensure that the age-related effects were not artifacts tied to individual samples, we averaged TE expression across brain regions for each individual and generated a PCA for the 22 subjects (Extended Data Fig. 3d).

In the brain disease dataset, RIN was found to have no effect on TE smRNAs and was excluded from the model. However, sequencing batches influenced the data structure, so we separated samples into two datasets based on the batches to avoid any bias, as not all batches had a representation of both Huntington’s and Parkinson’s samples. After data split, the batches were added as covariate in the DEA model. Post filtering very low expressed TEs for human data (mean <= 10 across all the samples/features) differential expression analysis was performed for aging between two age groups (0-20 yrs and 21-40 yrs for the human BrainSpan Atlas, 4 month and 24 month for mouse brains) and for disease status (HD or PD vs. control). Significance was corrected using Benjamini-Hochberg method, with a false discovery rate (FDR) threshold of < 0.01. Additionally, we applied a nested analysis of variance (AOV) method to account for individual effect (RIN corrected normalized expression ∼ brain region + age group / individual). After correcting for false discovery rate using Benjamini-Hochberg method, the P.adjust value was then compared to the DESeq2 results (Extended Data Fig. 3g).

Contrary to smRNA dataset, RIN showed no effect on the lnRNA dataset for both BrainSpan and aging mice and were not added as covariate in the DEA model. BrainSpan data was however corrected for batch effect. The lnRNA dataset for brain disease was also separated into two groups (HD and PD) based on the batch (Extended Data Fig. 8a), similar to smRNA. The data were corrected for batch as well as Age of Death (AOD) due to its effect on the clustering as seen in the PCA (Extended Data Figs. 8e, 9d) and on the distribution of the technical batches. PCA also highlighted two outliers in HD samples, confirmed by hierarchical clustering (Extended Data Fig. 8b, 8f), which were filtered out (sample number: 1104 and 1105). After filtering for very low expressed features (mean < 10 across all the samples/feature), differential expression analysis was performed on all the datasets for both TEs and genes separately using *DESeq2* in *R.* Since lnRNA BrainSpan data did not show a clear clustering like smRNA PCA even at further division of age groups (0-1yrs, 2-10yrs, 11-20yrs and 21-40yrs), same age groupings as for smRNA (0-20 yrs and 21-40 yrs) were chosen during DEA for consistency. Significance was corrected for false discovery rate using Benjamini-Hochberg method at FDR < 0.01.

### Correlation analyses and Intersection analyses between TE smRNAs and TE lnRNAs DE patterns

Normalized (log2 transformed) expression data of TE smRNAs and TE lnRNAs were matched based on overlapping samples for each of the four datasets (BrainSpan corrected for brain region, n=255; Mice, n=20; HD-Control corrected for batch, n=30; PD-Control corrected for batch, n=37) and r- and P-values of Pearson correlation was calculated for each TE sub-families. Significance was corrected for false discovery rate using Benjamini-Hochberg method at FDR < 0.01.

To find the overlapping significantly correlated genes and TE mRNAs we selected only significantly DE genes and TEs (FDR < 0.01) from human aging, HD and PD datasets. We performed correlation analysis on each of the datasets individually by calculating r-value of Pearson correlation between the genes and TEs corrected expression levels. Utilizing r-value > 0.5, significant gene-TE correlated pairs were filtered and an intersect analysis was performed to find the common gene-TE pairs across the three human datasets. From these shared pairs, we generated a network plot using *Cytoscape* ^96^ to demonstrate the relationship and to find common gene-TE biomarkers between aging and neurodegenerative diseases.

To find which dysregulated TEs were common across the human aging, HD and PD, we obtained the overlap of all TE smRNA and mRNA present in all human datasets. Using significance (FDR < 0.1) and log2fold change, we defined their up- or down-regulation in each dataset, leading to 6 possible groups. We then generated two UpSet plots in *R* to perform the intersect analysis separately for TE smRNAs and TE lnRNAs. To find the common TE biomarkers between TE smRNAs and TE lnRNAs, we first performed an intersect analysis between DE smRNA TE (FDR < 0.1) separately for aging-HD and aging - PD, irrespective of the directionality. We also did a similar intersect analysis for DE mRNA TEs (FDR < 0.1). We then intersected common TE smRNAs with common TE lnRNAs for aging-HD and aging-PD, using *Venn* diagram.

## Resource availability Lead contact

Further information and requests for resources and reagents should be directed to and will be fulfilled by the lead contact, Nelson Lau.

## Materials availability

All unique/stable reagents generated in this study are available from the lead contact. Material transfer agreements with Boston University may apply.

## Data and code availability

All sequencing data produced generated by this study is available on Sequencing Read Archive (SRA) under BioProject PRJNA1113634. See Tables S1 and S2 for specific BioSample and SRA accessions. SRA accessions for publicly available datasets used in this study can be found in Table S1C, S2C. The MSRG pipeline code can be found on the Github repository: https://github.com/laulabbumc/MosquitoSmallRNA

## ACKNOWLEDGEMENTS AND FUNDING

We thank members of the Lau lab for comments on this manuscript. N.C.L. is funded by NIH GRANT R01GM135215, R01AR078306, R01AG078930 and R01AG052465. B.K. CONNIZO is funded by R00AG063896 and R35GM151127. R.H.M. is funded by R01NS073947.

## Author contributions

Conceptualization, N.C.L, and G.D.; Methodology/Investigation, G.D., S.G., N.C.L.; Formal Analysis, G.D., S.G., and N.C.L.; Data Curation/Software, G.D. and S.G.; Sample Contribution & Resources, B.K.C. and R.H.M.; Writing – Original Draft, G.D. and N.C.L.; Writing – Review & Editing, G.D., S.G., B.K.C. R.H.M., A.T.L., B.C., and N.C.L.,; Visualization, G.D., A.T.L., and N.C.L.; Funding Acquisition, N.C.L., B.K.C., and R.H.M.;

---

**Extended Data Figure 1. Human BrainSpan Atlas smRNA compositions.**

(**a**) Barplot of the fractions of read counts broken down by all various smRNA categories per each BrainSpan Atlas sample. (**b**) Similar barplot of (**a**) but subtracting the major fraction of the miRNAs. (**c**) Boxplots of read counts comparison between pre-adults (0-20yrs) and adults (21-40yrs) brain samples for miRNAs, anti-sense TEs, sense TEs, piRNAs and structural RNAs. Statistical significance in the differences of these smRNA categories between young and aged brain samples is measured by ANOVA with p-values shown for each boxplot.

**Extended Data Figure 2. Mouse aging brains smRNA compositions.**

(**a**) Barplot of the fractions of read counts broken down by all various smRNA categories per each mouse brain sample. (**b**) Similar barplot of (**a**) but subtracting the major fraction of the miRNAs. (**c**) Boxplots of read counts comparison between young (4 months) and aged (24months) mouse brain samples for miRNAs, anti-sense TEs, sense TEs, piRNAs and structural RNAs. Statistical significance in the differences of these smRNA categories between young and aged brain samples is measured by ANOVA with p-values shown for each boxplot.

**Extended Data Figure 3. Human BrainSpan Atlas and mouse aging brains smRNA and lnRNA data processing.**

(**a-g**) Human BrainSpan Atlas data processing. PCA of smRNA based on RNA Integrity Number (RIN, **a**) or Cause of Death (COD, **b**). (**c**) Correlation between various smRNA associated metadata. (**d**) PCA for averaged counts across brain regions for 22 individuals in the BrainSpan Atlas. (**e**) Differentially expressed TE smRNAs between pre-adults and adults. (**f**) Scatterplot of P-adjusted values from nested analysis of variance (AOV) compared to DESeq2 results. (**g**) Differentially expressed human BrainSpan Atlas TE lnRNAs between pre-adult (0-20 yrs) and adults (21-40 yrs) at FDR < 0.01. (**h-l**) Aging mouse brains data processing. (**h**) PCA of smRNA based on RIN. (**i**) Differentially expressed TE smRNAs between young (4months) and aged mice (24months) at FDR < 0.01. (**j**) Differentially expressed TE smRNAs between young (4months) and aged mice (24months) at P-value < 0.05. (**k**) Differentially expressed mouse aging brains TE lnRNAs between young and aged mice at FDR < 0.01 (**l**) Differentially expressed mouse aging lnRNA genes and TEs between young and aged mice at FDR < 0.01.

**Extended Data Figure 4. Human BrainSpan Alu smRNA coverage, human brain TE RNA clustering, human and mouse aging brains lnRNA gene clustering.**

(**a**) Mean coverage plot of human aging TE smRNA across Alu consensus TE across all samples in pre-adults (0-20 yrs) (**b**) Mean coverage plot of human aging TE smRNA across Alu consensus TE across all samples in adults (21-40 yrs). (**c**) Heatmap clustering based on z-score of the top highly expressed TEs across brain regions. (**d**) Heatmap clustering for top human brain lnRNA DE genes between pre-adults (0-20 yrs) and adults (21-40 yrs). (**e**) Heatmap clustering for top mouse brain lnRNA DE genes between young (4 months) and aged (24 months).

**Extended Data Figure 5. Human brain smRNA compositions in the HD-Control and PD-Control cohorts.**

(**a**) Barplot of the fractions of read counts broken down by all various smRNA categories in HD, PD and control cohorts samples. (**b**) Similar barplot of (**a**) but subtracting the major fraction of the miRNAs.

**Extended Data Figure 6. Differential Expression Analysis (DEA) of human brain TE smRNA in HD-Control and PD-Control cohorts.**

(**a**) Boxplots of read count comparison of miRNA, structural RNAs, anti-sense TEs, sense TEs, and piRNAs between HD-vs-Controls cohorts. (**b**) Volcano plot of differentially expressed smRNA TEs between HD-control at FDR < 0.01. (**c**) Boxplots of read count comparison of miRNA, structural RNAs, anti-sense TEs, sense TEs, and piRNAs between PD-vs-Controls cohorts. (**d**) Volcano plot of differentially expressed smRNA TEs between PD-control at FDR < 0.05.

**Extended Data Figure 7. HD and PD TE smRNA characteristics.**

(**a**) Length distribution of smRNA TEs compared in Huntington’s, Parkinson’s and Controls brain samples. (**b**) Sequence logos of the base compositions in smRNA reads of Huntington’s, Parkinson’s and Controls brain samples.

**Extended Data Figure 8. Human brain lnRNA characteristics and outliers’ analysis in the HD-Control and PD-Control cohorts.**

(**a**) Boxplot of Age Of Death (AOD) frequencies across the sample batches. (**b**) PCA showing batch effect on PD and control data based off the lnRNA data. (**c**) Boxplot of AOD frequencies showing a correlation with BRAAK scores. (**d**) PCA showing two outliers for HD-control data pointed to by the hashed box. (**e**) Heatmap clustering showing altered TE expression structure for the two outlier HD samples also pointed to by the hashed box. (**f**) PCA showing AOD effect on HD and control cohorts from the lnRNA data after outlier removal.

**Extended Data Figure 9. Human brain lnRNA Differentially Expressed TEs and Genes in the HD-Control and PD-Control cohorts.**

(**a**) Differentially expressed TE lnRNAs between PD-control at FDR <0.01. (**b**) Differentially expressed TE lnRNAs between HD-control at FDR <0.0. (**c**) PCA showing clustering of all features (genes and TEs) based on disease status (HD, PD, control). (**d**) PCA showing effect of AOD on all features (genes and TEs). (**e**) PCA showing clustering of all features (genes and TEs) based on disease status for HD-vs-Control. (**f**) PCA showing clustering of all features (genes and TEs) based on disease status for PD-vs-Control. (**g**) Differentially expressed genes and TEs lnRNAs between HD-vs-Control at FDR < 0.01. (**h**) Differentially expressed genes and TEs lnRNAs between HD-vs-Control at FDR < 0.01.

**Extended Data Figure 10. Gene Set Enrichment analysis of the lnRNAs from the BrainSpan Atlas and the the HD-Control and PD-Control cohorts.**

(**a**) Significantly enriched pathways against BioCarta using overlapping genes between aging, HD and PD with FDR cutoff of 0.05. (**b**) Significantly enriched pathways against KEGG using overlapping genes between aging, HD and PD with FDR cutoff of 0.05.

**Extended Data Table 1. Metadata tabs of the BrainSpan Atlas smRNA and lnRNA and mouse brain smRNA and lnRNA sequencing datasets.**

**Extended Data Table 2. Count matrix tabs of the BrainSpan Atlas TE smRNAs and TE lnRNAs Differential Expression (DE) and biomarker candidate highlights.**

**Extended Data Table 3. Count matrix tabs of the aging mouse brains TE smRNAs and TE lnRNAs Differential Expression (DE).**

**Extended Data Table 4. Metadata tabs of the PD-Control and HD-Control brain smRNA and lnRNA datasets, adapted from**^62,64,65,73^.

**Extended Data Table 5. Count matrix tabs of the PD-Control and HD-Control TE smRNAs and TE lnRNAs Differential Expression (DE) and biomarker candidate highlights.**

**Extended Data Table 6. Correlation values from the intersection analyses of gene versus TE DE patterns between BrainSpan aging and HD and PD.**

