## Supplemental Figure S1 for "Transposable element small and long RNAs in aging brains and implications in Huntington’s and Parkinson’s disease"

Dayama, et al.
Extended Data Figure 1.

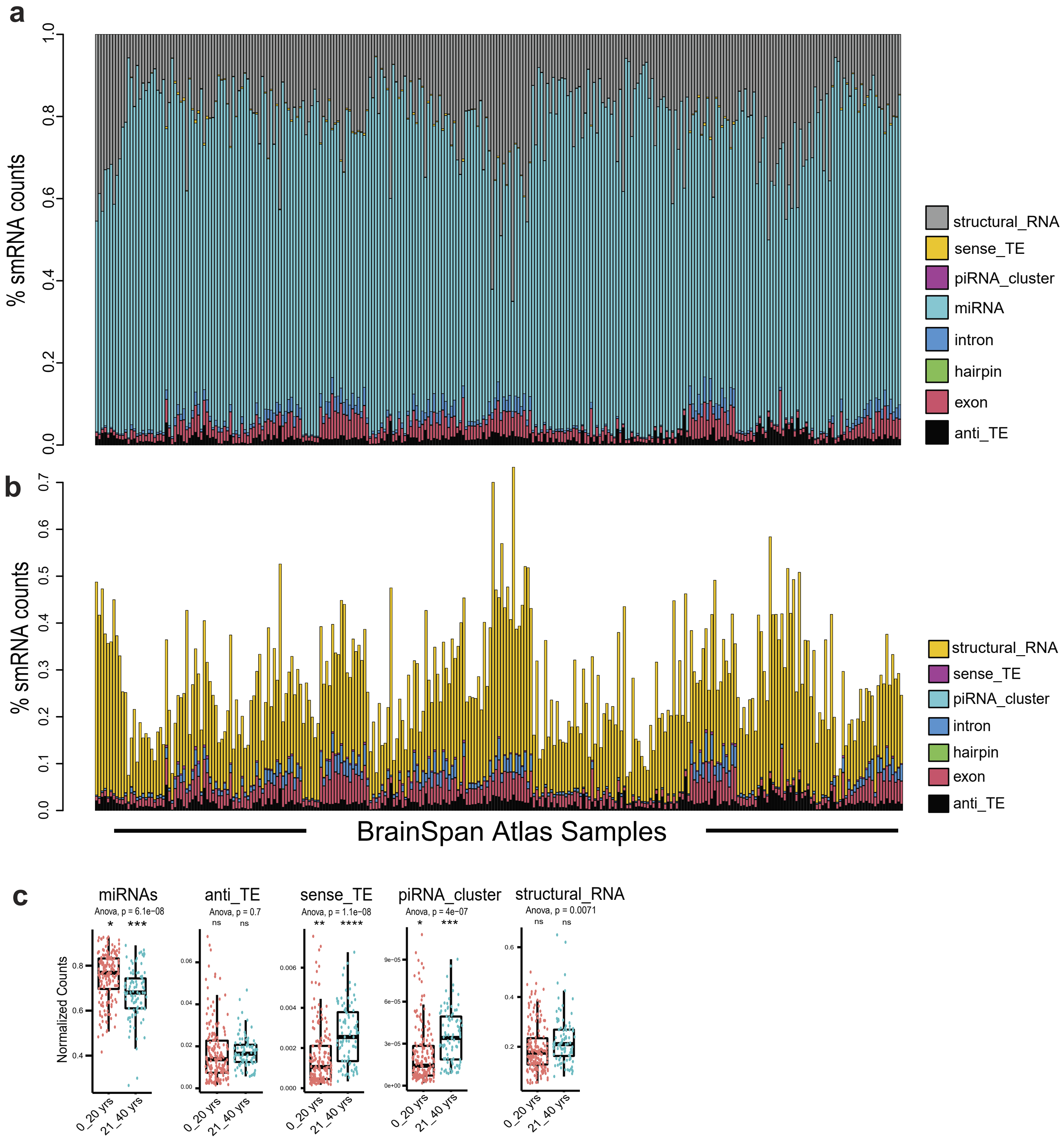


**Extended Data Figure 1. Human BrainSpan Atlas smRNA compositions.**
(**a**) Barplot of the fractions of read counts broken down by all various smRNA categories per each BrainSpan Atlas sample. (**b**) Similar barplot of (A) but subtracting the major fraction of the miRNAs. (**c**) Boxplots of read counts comparison between pre-adults (0-20yrs) and adults (21-40yrs) brain samples for miRNAs, anti-sense TEs, sense TEs, piRNAs and structural RNAs. Statistical significance in the differences of these smRNA categories between young and aged brain samples is measured by ANOVA with p-values shown for each boxplot.
