## Supplemental Figure S2 for "Transposable element small and long RNAs in aging brains and implications in Huntington’s and Parkinson’s disease"

Dayama, et al.
Extended Data Figure 2.

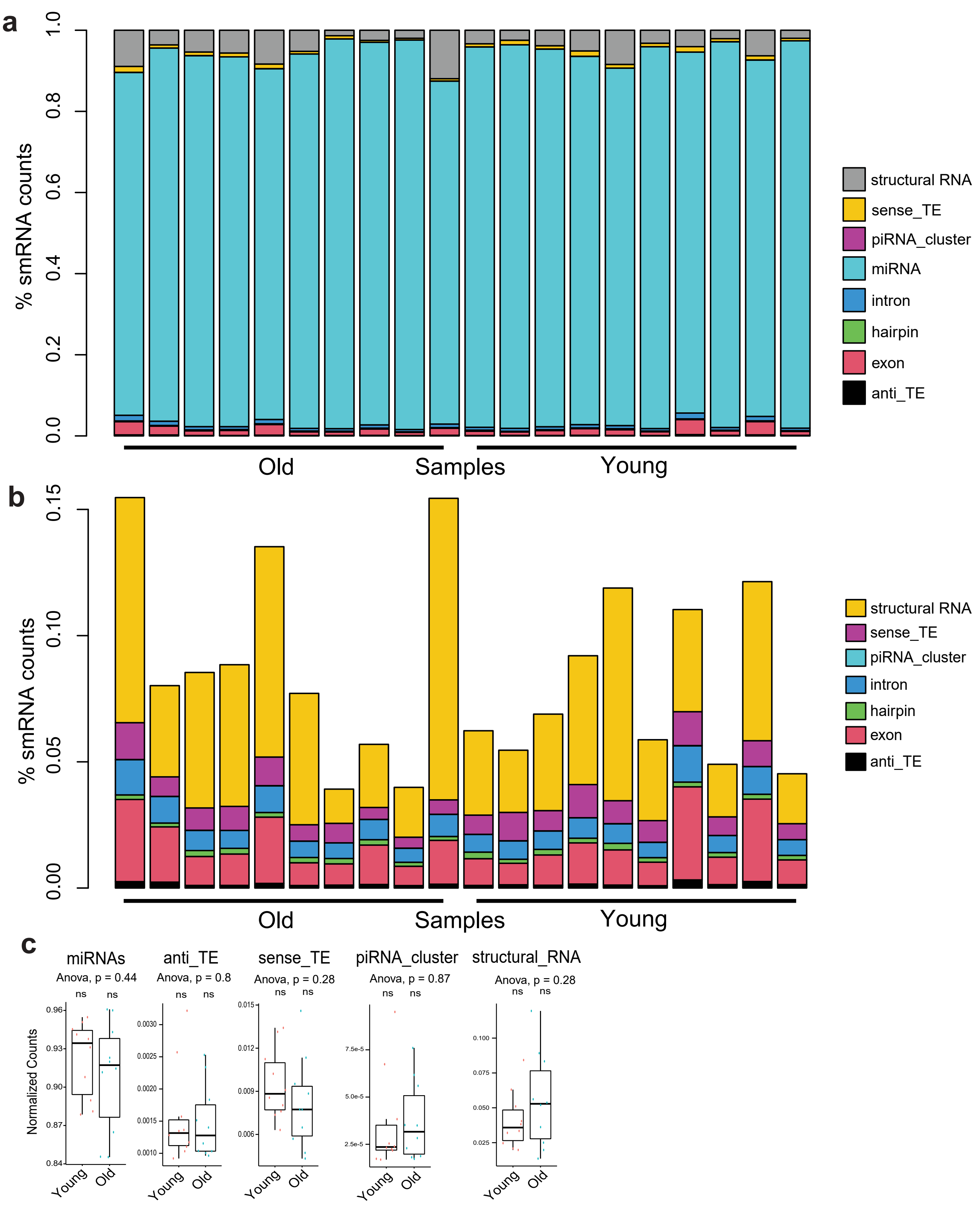


**Extended Data Figure 2. Mouse aging brains smRNA compositions.**
(**a**) Barplot of the fractions of read counts broken down by all various smRNA categories per each mouse brain sample. (**b**) Similar barplot of (**a**) but subtracting the major fraction of the miRNAs. (**c**) Boxplots of read counts comparison between young (4 months) and aged (24months) mouse brain samples for miRNAs, anti-sense TEs, sense TEs, piRNAs and structural RNAs. Statistical significance in the differences of these smRNA categories between young and aged brain samples is measured by ANOVA with p-values shown for each boxplot.
