## Supplemental Figure S3 for "Transposable element small and long RNAs in aging brains and implications in Huntington’s and Parkinson’s disease"

Dayama et al.
Extended Data Figure 3.

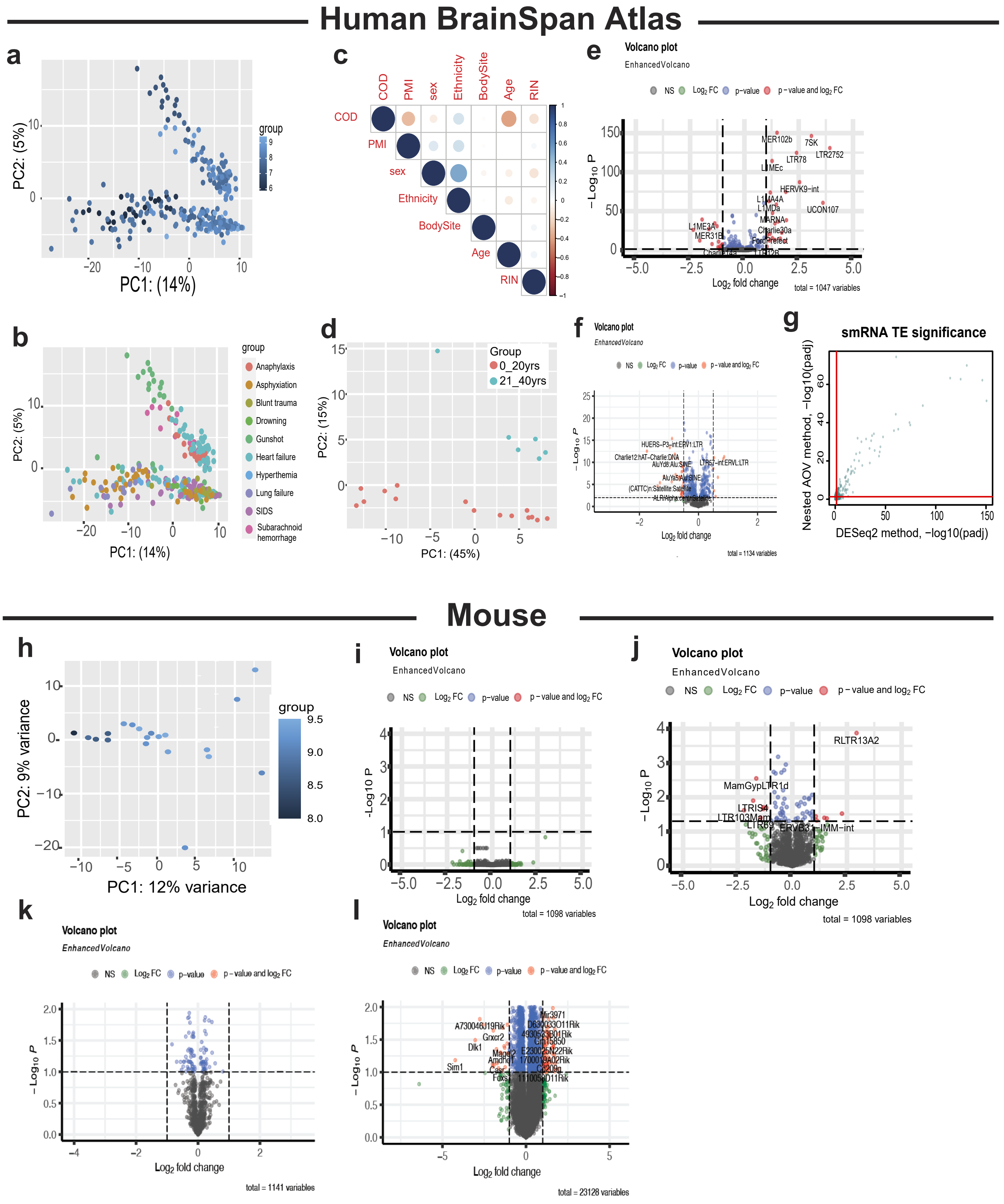


**Extended Data Figure 3. Human BrainSpan Atlas and mouse aging brains smRNA and lnRNA data processing.** (**a-g**) Human BrainSpan Atlas data processing. PCA of smRNA based on RNA Integrity Number (RIN, **a**) or Cause of Death (COD, **b**). (**c**) Correlation between various smRNA associated metadata. (**d**) PCA for averaged counts across brain regions for 22 individuals in the Brain-Span Atlas. (**e**) Differentially expressed TE smRNAs between pre-adults and adults. (**f**) Scatterplot of P-adjusted values from nested analysis of variance (AOV) compared to DESeq2 results. (**g**) Differentially expressed human BrainSpan Atlas TE lnRNAs between pre-adult (0-20 yrs) and adults (21-40 yrs) at FDR < 0.01. (**h-l**) Aging mouse brains data processing. (**h**) PCA of smRNA based on RIN. (**i**) Differentially expressed TE smRNAs between young (4months) and aged mice (24months) at FDR < 0.01. (**j**) Differentially expressed TE smRNAs between young (4months) and aged mice (24months) at P-value < 0.05. (**k**) Differentially expressed mouse aging brains TE lnRNAs between young and aged mice at FDR < 0.01. (**l**) Differentially expressed mouse aging lnRNA genes and TEs between young and aged mice at FDR < 0.01.
