## Supplemental Figure S4 for "Transposable element small and long RNAs in aging brains and implications in Huntington’s and Parkinson’s disease"

**Dayama et al.**
**Extended Data Figure 4.**

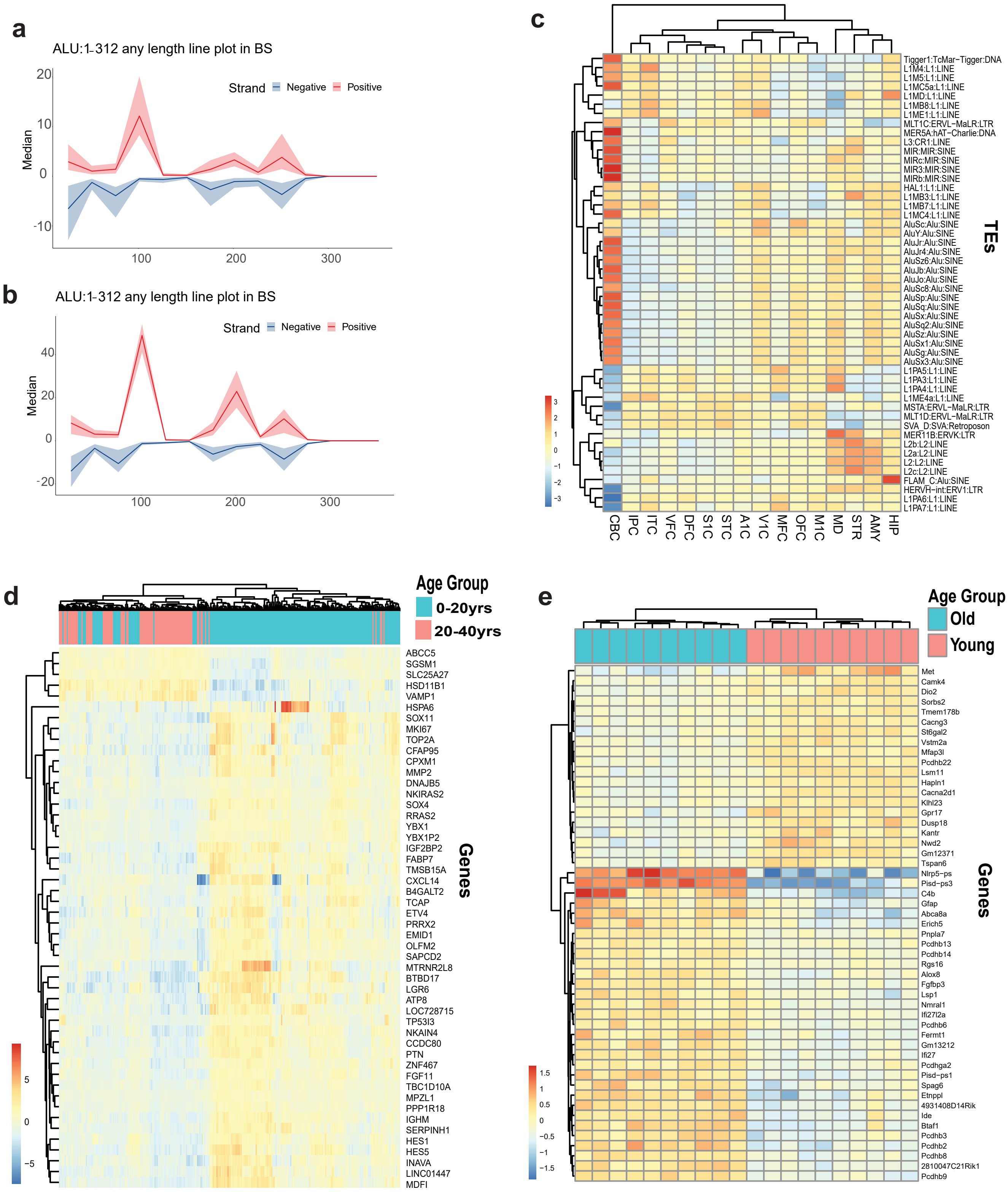


**Extended Data Figure 4. Human BrainSpan Alu smRNA coverage, human brain TE RNA clustering, human and mouse aging brains lnRNA gene clustering.**
(**a**) Mean coverage plot of human aging TE smRNA across Alu consensus TE across all samples in pre-adults (0-20 yrs). (**b**) Mean coverage plot of human aging TE smRNA across Alu consensus TE across all samples in adults (21-40 yrs). (**c**) Heat-map clustering based on z-score of the top highly expressed TEs across brain regions. (**d**) Heatmap clustering for top human brain lnRNA DE genes between pre-adults (0-20 yrs) and adults (21-40 yrs). (**e**) Heatmap clustering for top mouse brain lnRNA DE genes between young (4 months) and aged (24 months).
