## Supplemental Figure S5 for "Transposable element small and long RNAs in aging brains and implications in Huntington’s and Parkinson’s disease"

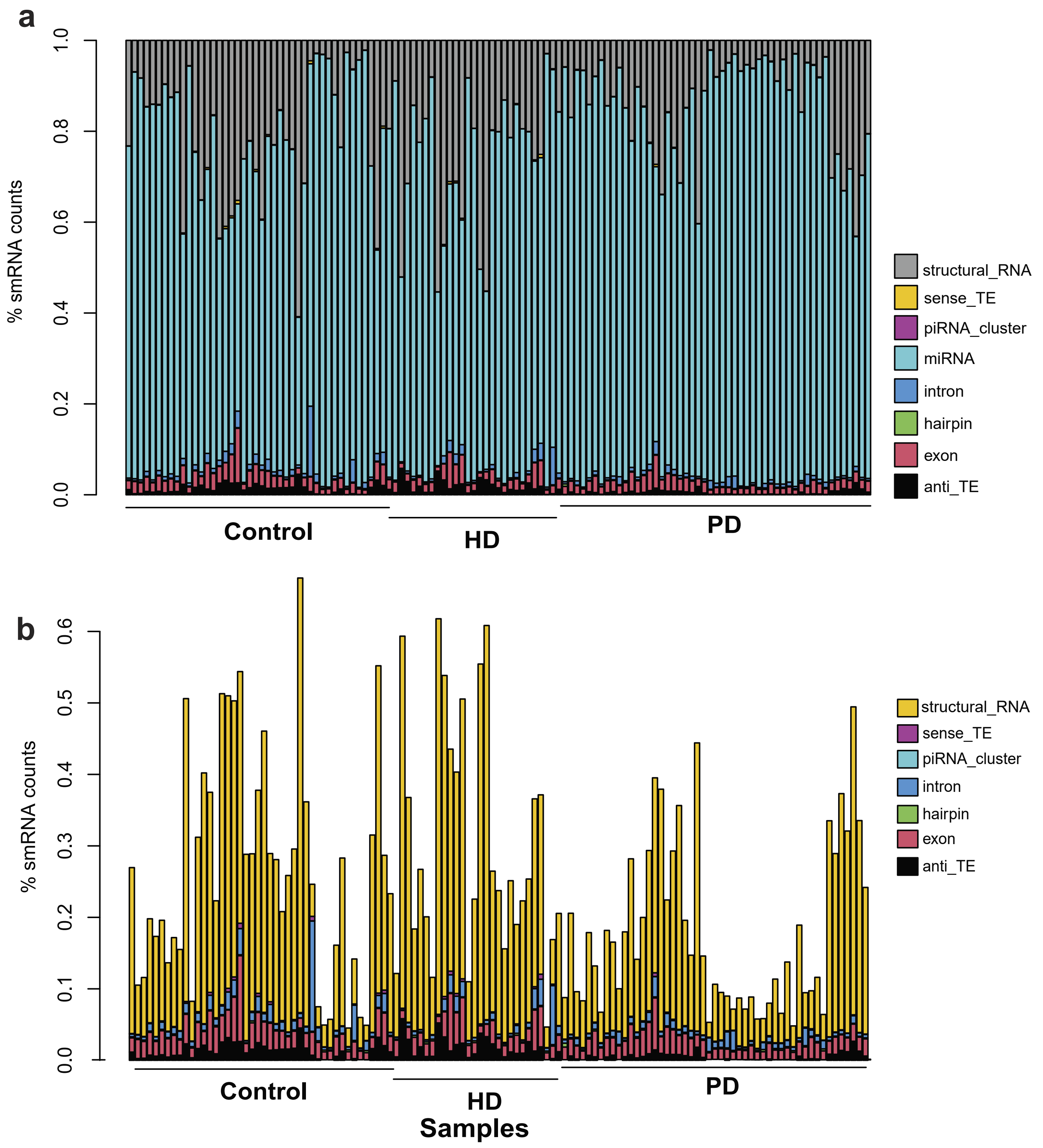


**Extended Data Figure 5. Human brain smRNA compositions in the HD-Control and PD-Control cohorts.**
(**a**) Barplot of the fractions of read counts broken down by all various smRNA categories in the HD, PD and control cohorts samples. (**b**) Similar barplot of (**a**) but subtracting the major fraction of the miRNAs.
