## Supplemental Figure S6 for "Transposable element small and long RNAs in aging brains and implications in Huntington’s and Parkinson’s disease"

Dayama et al.
Extended Data Figure 6.

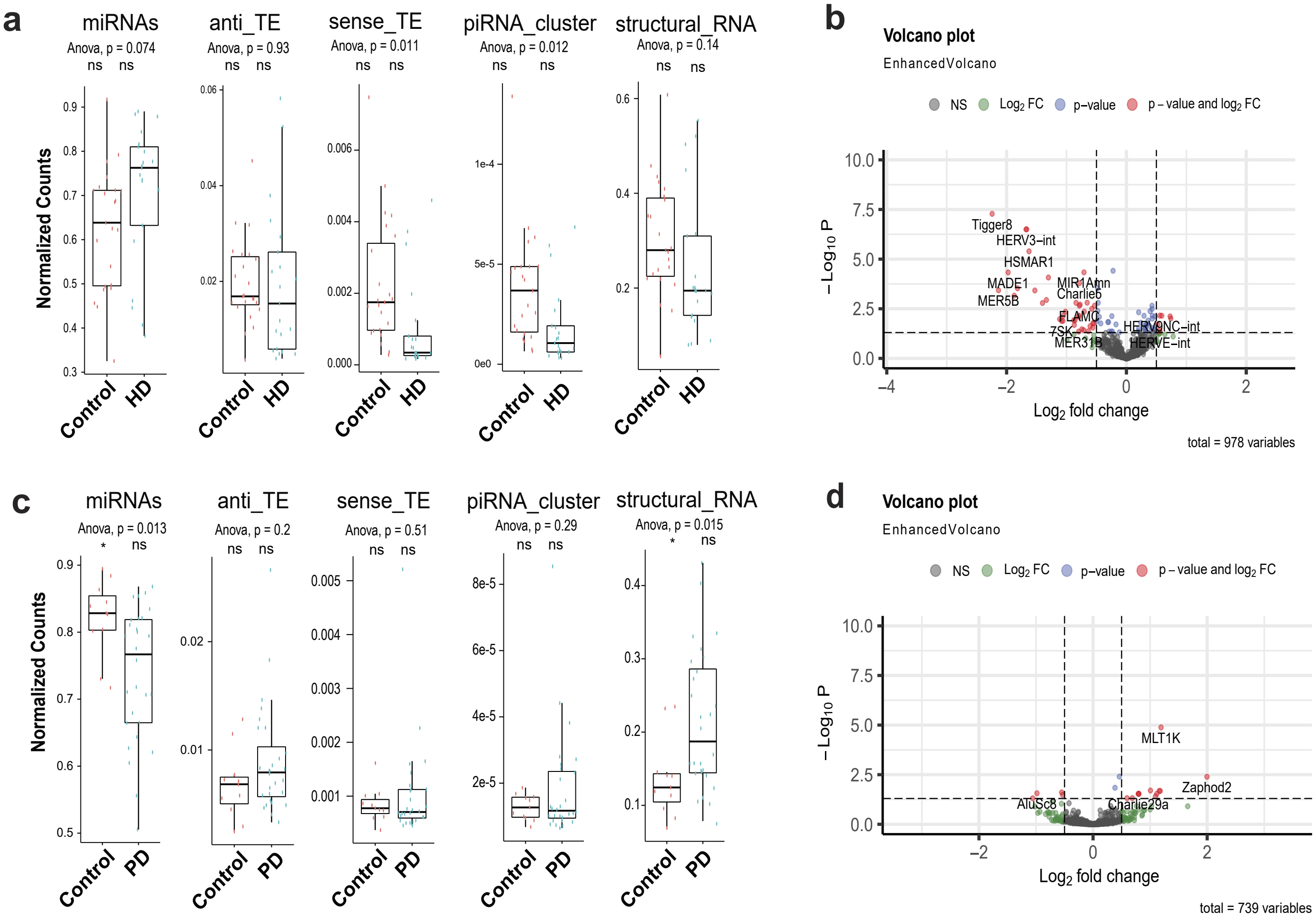


**Extended Data Figure 6. Differential Expression Analysis (DEA) of human brain TE smRNA in HD-Control and PD-Control cohorts.**
(**a**) Boxplots of read count comparison of miRNA, structural RNAs, anti-sense TEs, sense TEs, and piRNAs between HD-vs-Controls cohorts. (**b**) Volcano plot of differentially expressed smRNA TEs between HD-control at FDR < 0.01. (**c**) Boxplots of read count comparison of miRNA, structural RNAs, anti-sense TEs, sense TEs, and piRNAs between PD-vs-Controls cohorts. (**d**) Volcano plot of differentially expressed smRNA TEs between PD-control at FDR < 0.05.
