## Supplemental Figure S7 for "Transposable element small and long RNAs in aging brains and implications in Huntington’s and Parkinson’s disease"

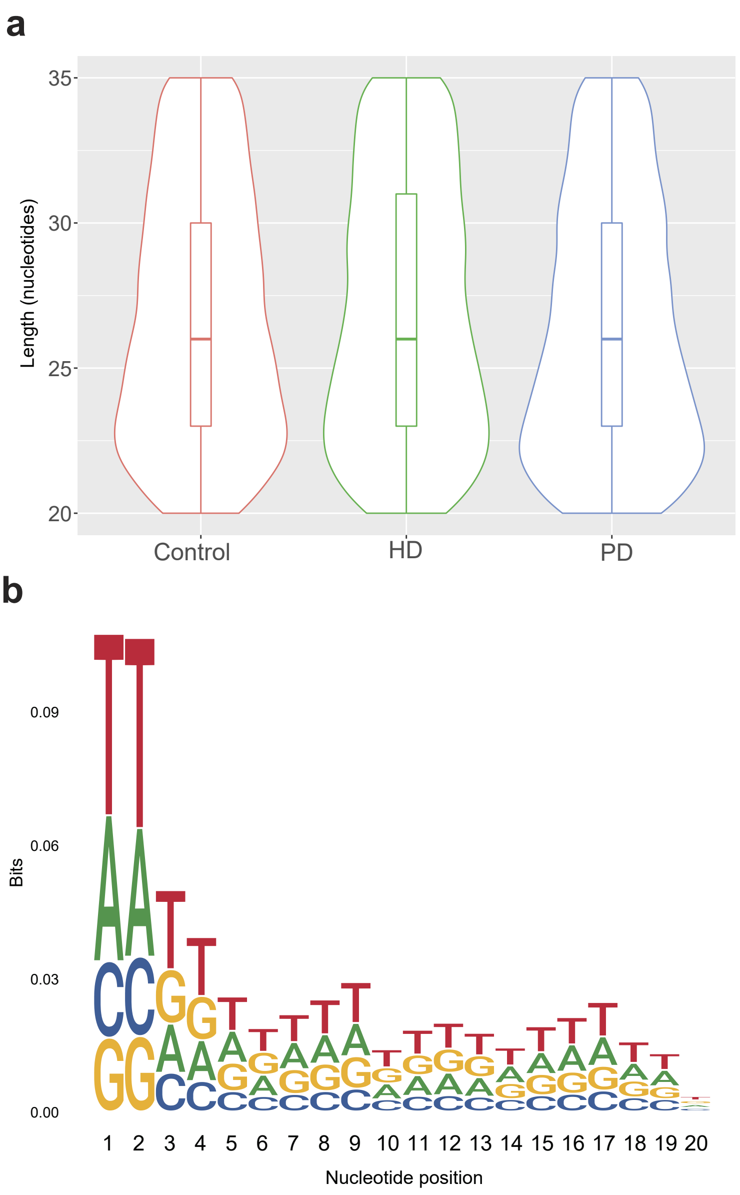


**Extended Data Figure 7. HD and PD smRNA characteristics.**
(**a**) Length distribution of smRNA TEs compared in Huntington's, Parkinson's and Controls brain samples. (**b**) Sequence logos of the base compositions in smRNA reads of Huntington's, Parkinson's and Controls brain samples.
