## Supplemental Figure S8 for "Transposable element small and long RNAs in aging brains and implications in Huntington’s and Parkinson’s disease"

Dayama et al.
Extended Data Figure 8.

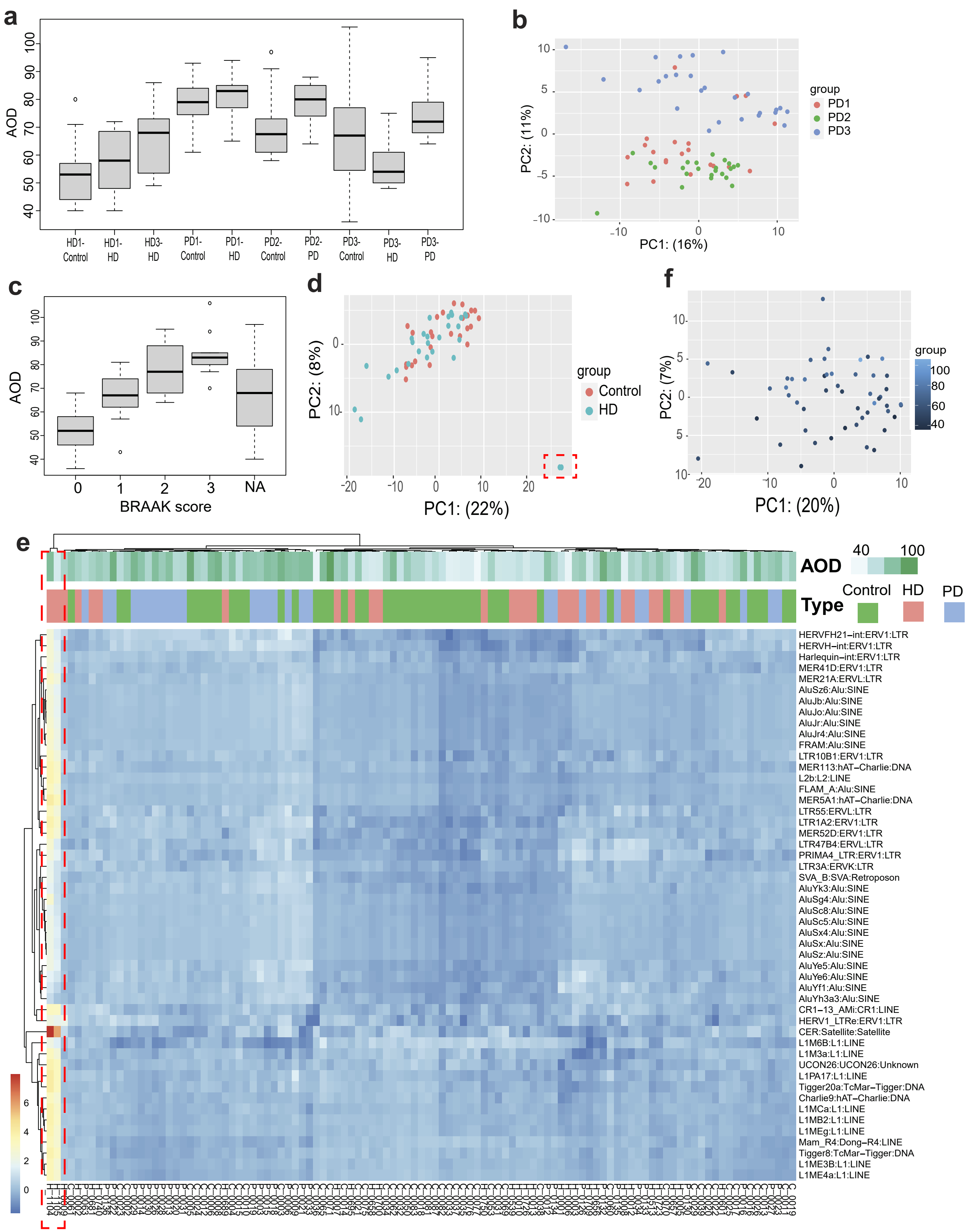


**Extended Data Figure 8. Human brain lnRNA characteristics and outliers' analysis in the HD-Control and PD-Control cohorts.**
(a) Boxplot of Age Of Death (AOD) frequencies across the sample batches. (b) PCA showing batch effect on PD and control data based off the lnRNA data. (c) Boxplot of AOD frequencies showing a correlation with BRAAK scores. (d) PCA showing two outliers for HD-control data pointed to by the hashed box. (e) Heatmap clustering showing altered TE expression structure for the two outlier HD samples also pointed to by the hashed box. (f) PCA showing AOD effect on HD and control cohorts from the lnRNA data after outlier removal.
