## Supplemental Figure S9 for "Transposable element small and long RNAs in aging brains and implications in Huntington’s and Parkinson’s disease"

Dayama et al.
Extended Data Figure 9.

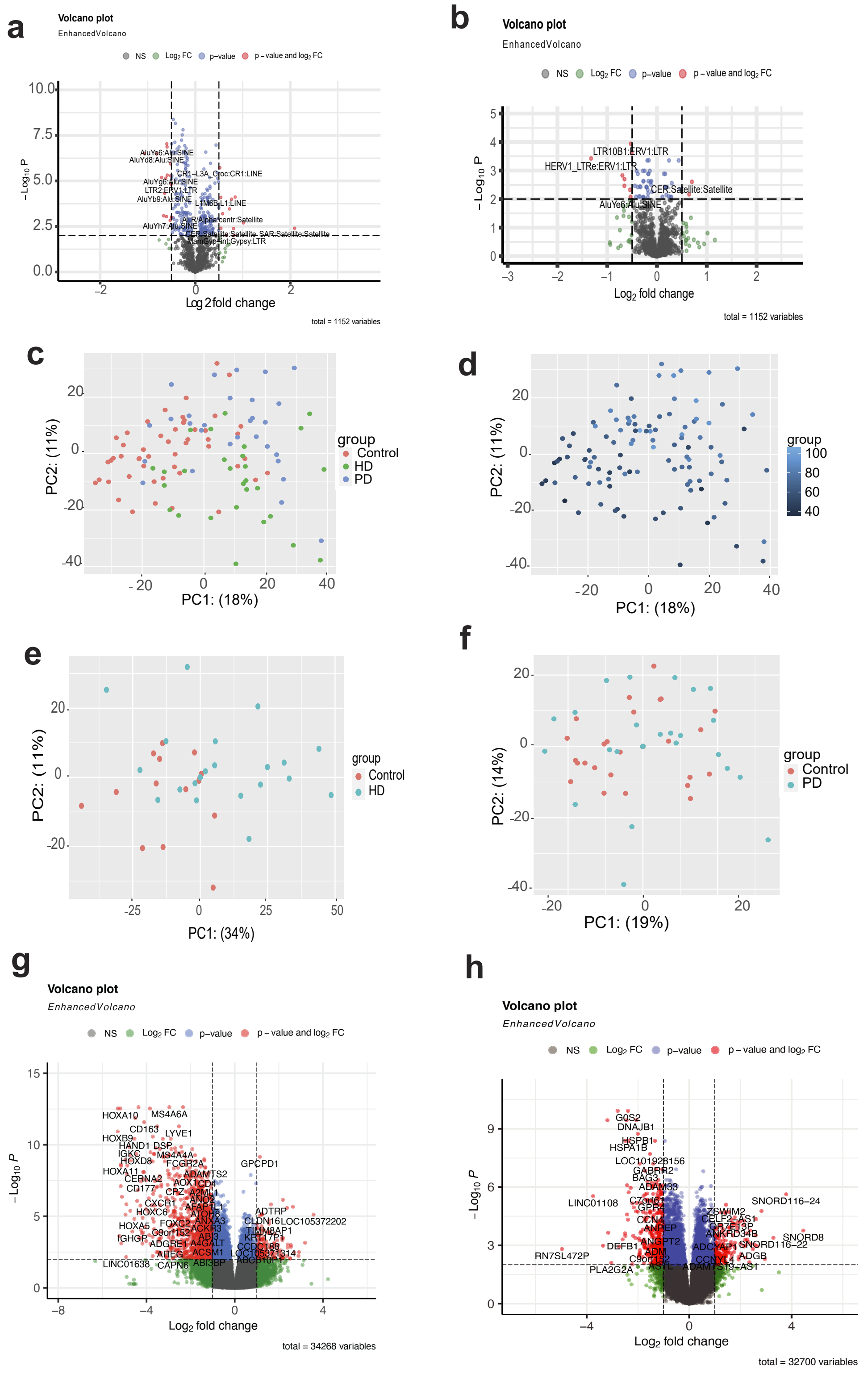

**Extended Data Figure 9. Human brain lnRNA Differentially Expressed TEs and Genes in the HD-Control and PD-Control cohorts.**
(**a**) Differentially expressed TE lnRNAs between PD-control at FDR <0.01. (**b**) Differentially expressed TE lnRNAs between HD-control at FDR <0.0. (**c**) PCA showing clustering of all features (genes and TEs) based on disease status (HD, PD, control). (**d**) PCA showing effect of AOD on all features (genes and TEs). (**e**) PCA showing clustering of all features (genes and TEs) based on disease status for HD-vs-Control. (**f**) PCA showing clustering of all features (genes and TEs) based on disease status for PD-vs-Control. (**g**) Differentially expressed genes and TEs lnRNAs between HD-vs-Control at FDR < 0.01. (**h**) Differentially expressed genes and TEs lnRNAs between HD-vs-Control at FDR < 0.01.
