## Supplemental Figure S10 for "Transposable element small and long RNAs in aging brains and implications in Huntington’s and Parkinson’s disease"

**Dayama et al.**
**Extended Data Figure 10.**

**a**

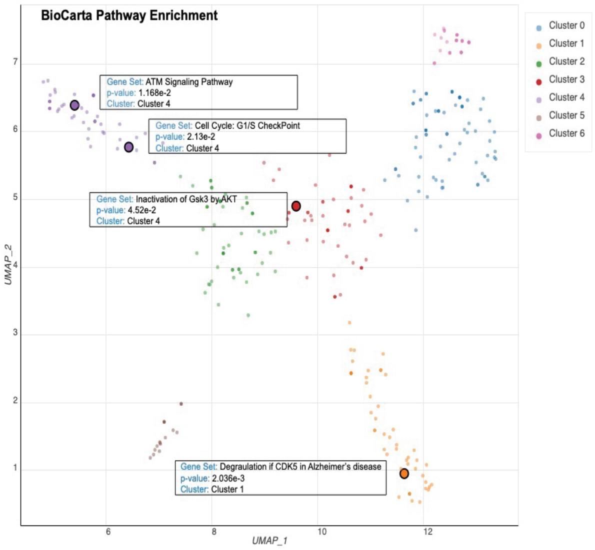


**b**

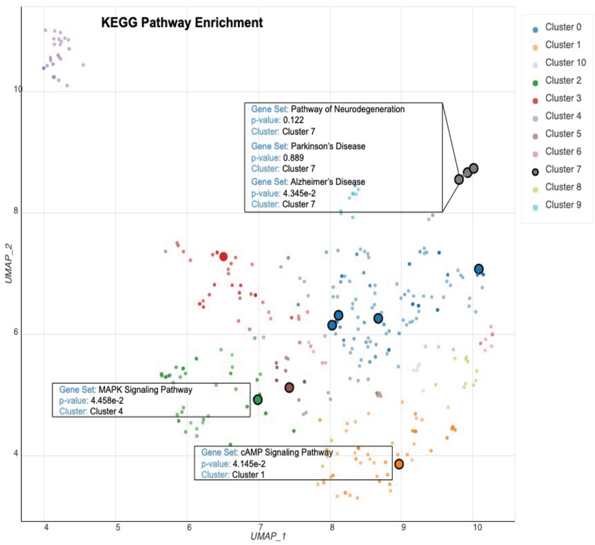


**Extended Data Figure 10. Gene Set Enrichment analysis of the lnRNAs from the BrainSpan Atlas and the the HD-Control and PD-Control cohorts.**
(**a**) Significantly enriched pathways against BioCarta using overlapping genes between aging, HD and PD with FDR cutoff of 0.05. (**b**) Significantly enriched pathways against KEGG using overlapping genes between aging, HD and PD with FDR cutoff of 0.05.
